# Collateral Immune Cell Signaling Compromises the Efficacy of T-cell Engaging Therapies for Cardiac Fibrosis

**DOI:** 10.64898/2026.09.15.751626

**Authors:** Steven Yang, Alekhya Parvathaneni, Yun-Ling Pai, Andrew Koenig, Farid Kadyrov, Macee C Owen, Junedh M Amrute, Andrea Bredemeyer, Yu-Sung Hsu, Attila Kovacs, Carla J Weinheimer, Jessica M Nigro, Lance Yeh, Thomas Williamson, Blake Jardin, Joel G Rurik, Melissa Thomas, Jinghong Wang, Shweta Mandavalli, Kevin Cook, Tracy Yamawaki, Shibali Das, Niklas Hartmann, Laura M Wienecke, Candice Baker, Chi-Ming Li, Florian Leuschner, Nadia Rosenthal, Jonathan A Epstein, Nathan Singh, Simon Jackson, Brandon Ason, Kory J Lavine

## Abstract

**Cardiac fibrosis is causally linked to heart failure progression and survival. Currently, there are no approved treatments that directly target cardiac fibrosis. Recent studies have identified a subset of activated cardiac fibroblasts distinct from myofibroblasts that are marked by fibroblast activation protein (FAP) expression, emerge in the injured and diseased heart through inflammatory signaling, and contribute to fibrosis. Using a genetic mouse model, we demonstrate the potential benefits of FAP^+^ fibroblast depletion following myocardial infarction. Unexpectedly, while FAP targeted bispecific T-cell engaging antibodies (BiTE^®^ molecules) effectively eliminate FAP^+^ fibroblasts from the heart, they surprisingly lead to accelerated deterioration of cardiac function, enhanced remodeling, and increased scar size. FAP BiTE^®^ molecules elicit a robust cytokine response within the heart with prominent activation of interferon gamma (IFNγ) and CD40 ligand pathways. Target cell killing was independent of IFNγ and CD40L signaling and blockade of these pathways was sufficient to unmask the protective therapeutic effects of FAP^+^ fibroblast depletion. Mechanistically, we reveal that IFNγ signaling to fibroblasts drives the differentiation of an independent lineage of activated fibroblasts not typically found in the infarcted heart, which are responsible for the harmful effects of FAP BiTE^®^ molecules. Collectively, these findings highlight a previously unrecognized cardiac liability of BiTE^®^ molecules and inform the design of the next generation of therapeutics.**

## Introduction

Myocardial fibrosis impairs compliance of the heart and compromises cardiac function in both systole and diastole. Fibroblasts within the heart are activated in response to tissue injury and disease and are responsible for deposition of collagen into the extracellular matrix. Although fibrosis may help maintain myocardial integrity in select scenarios, chronic tissue injury and hemodynamic load stimulate excessive collagen deposition, which adversely affects ventricular function and drives heart failure^1,2^. Across numerous clinical studies, cardiac fibrosis is associated with worsening heart failure and increased frequency of adverse cardiovascular events^3,4^. Unfortunately, there are few therapeutics available to prevent or reduce cardiac fibroblast activation and tissue fibrosis. Development of such therapies for cardiac applications has remained particularly challenging^5,6^.

Advancements in sequencing technology have unveiled unanticipated heterogeneity amongst activated fibroblasts within the human heart. Several groups have revealed that fibroblast activation protein (FAP) is expressed on the cell surface of an activated fibroblast subset that expands following cardiac injury and is distinct from myofibroblasts^7–9^. Identification of the FAP^+^ fibroblast cell state has created new opportunities to selectively target specific subsets of activated fibroblasts to mitigate cardiac fibrosis. Consistent with this concept, CAR T cells targeting FAP^+^ fibroblasts reduce cardiac fibrosis in mouse models^7,10^. Given the complexity of CAR T cell production, there is incredible interest in developing off-the-shelf therapies to accelerate translation. Here, we investigate the utility of bispecific T-cell engaging antibodies for *in vivo* removal of FAP^+^ cardiac fibroblasts.

## Results

### Genetic ablation of *Fap^+^* specified fibroblasts improves cardiac function after injury

To study *Fap^+^* fibroblasts following cardiac injury, we have generated knock-in mice bearing a tamoxifen-inducible Cre under control of the native *Fap* regulatory elements (**Fig. 1A**). To model cardiac ischemic reperfusion injury (IRI), we utilized a newly developed minimally invasive approach that circumvents the need for surgical thoracotomy and thus minimizes confounding inflammatory and fibrotic responses related to instrumentation^11^ (**fig. S1**). We observed virtually no tdTomato^+^ cells present in the myocardium when uninjured *Fap^ert2Cre^ Rosa26^Lox-STOP-Lox-tdTomato^*(Fap tdT) reporter mice were administered daily tamoxifen for 1 week. However, following IRI, there was a robust increase in the number of tdTomato^+^ cells predominantly within the infarct. Consistent with studies in other injury models^12–14^, tdTomato^+^ cells were present by day 8 after IRI and persisted for at least 28 days thereafter (**Fig. 1B**). These data demonstrate that *Fap^+^* fibroblasts are found exclusively in the injured heart, are specified early following cardiac IRI, and this lineage of fibroblasts persists during more chronic stages of post-infarct remodeling.

**Figure 1.**
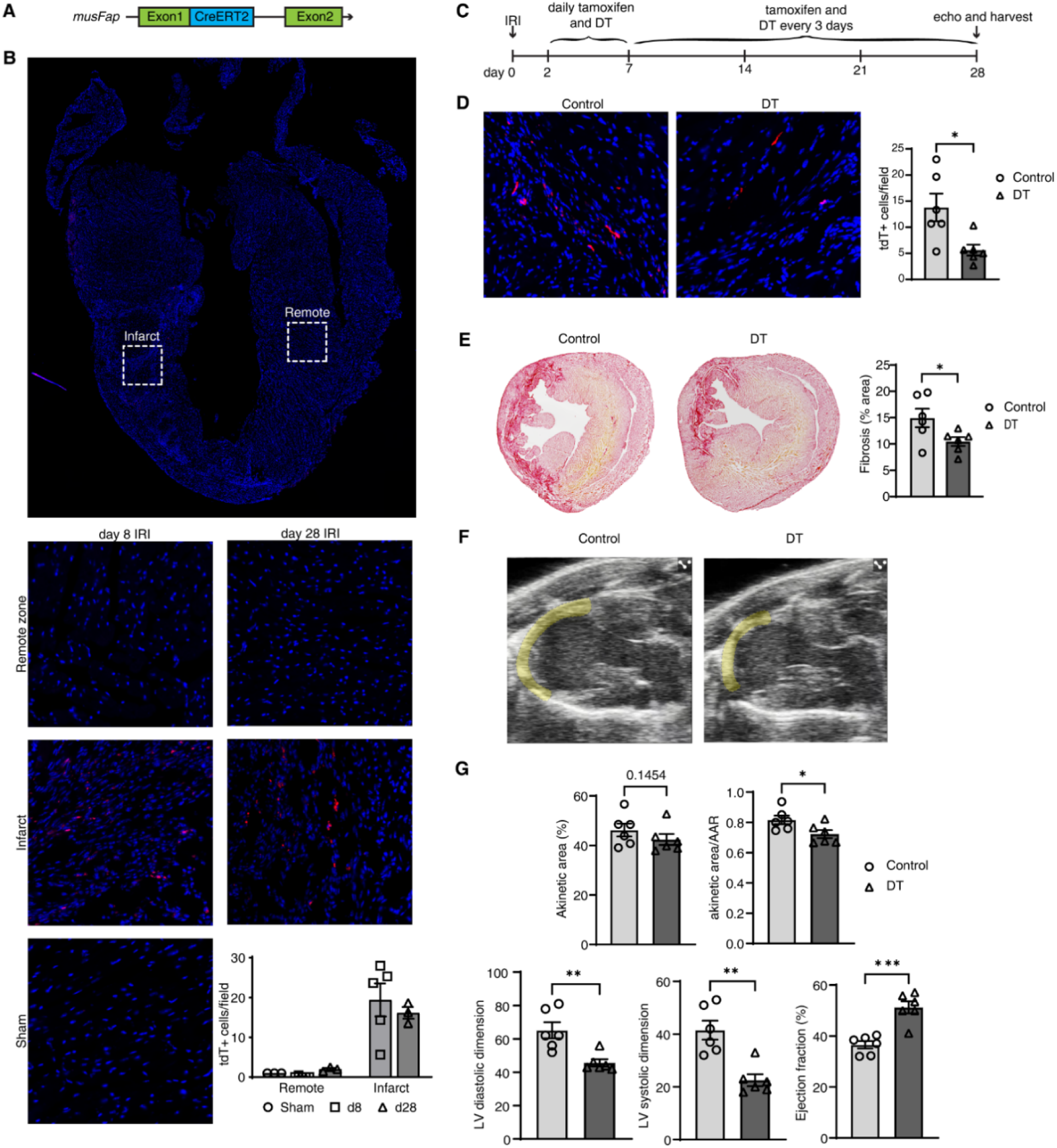
Ischemia reperfusion injury increases FAP^+^ fibroblasts and depletion of these cells ameliorates adverse chamber remodeling, improves systolic function, and decreases scar size after IRI. A) Schematic of *Fap^ert2Cre^*mouse B) tdTomato expression in *Fap* lineage reporter mouse without injury, and 8 or 28 days post IRI in the remote zone and infarct C) Experimental schematic D) tdTomato expression and quantification in control and DT treated Fap tdT DTR mice after IRI E) Picro Sirius Red staining and quantification F) Representative echocardiogram stills in systole with akinetic area highlighted G) Echocardiogram quantification

Despite growing interest in FAP as a therapeutic target, it has not been determined whether genetic depletion of Fap^+^ cells improves outcomes following cardiac injury. Fap tdT reporter mice were crossed to *Rosa26^Lox-STOP-Lox-DTR^* mice to generate a mouse line (Fap tdT DTR) to study the effects of ablating *Fap^+^* cells and their progeny with diphtheria toxin (DT). We dosed Fap tdT DTR mice with daily tamoxifen and DT or PBS after IRI (**Fig. 1C**). Immunofluorescence of Fap tdT DTR mice revealed a decrease in the number of tdTomato^+^ cells within the infarct in DT treated mice compared to PBS treated controls (**Fig. 1D**). Picrosirius red staining on day 28 after IRI revealed reduced fibrosis and smaller infarct size in DT treated mice (**Fig. 1E**). Echocardiography performed on day 28 after IRI demonstrated that depletion of *Fap^+^* cells reduced akinetic area, markedly diminished LV chamber dilation, and increased ejection fraction (**Fig. 1F-G**). No evidence of LV rupture or mortality was observed. These findings establish proof-of-concept that targeted depletion of Fap^+^ fibroblasts after cardiac injury may represent a viable therapeutic strategy.

### FAP targeted T cell engaging antibodies promote adverse remodeling and deteriorate cardiac function after injury

Having demonstrated that removing *Fap^+^* fibroblasts and their progeny is sufficient to mitigate remodeling and improve cardiac function following IRI, we sought to investigate an off-the-shelf and clinically tractable platform for FAP cell depletion, FAP targeted bispecific T cell engagers (BiTE^®^ molecules). This system builds off the success of CAR T cell technologies, but avoids challenges associated with cell product engineering for individual patients. Similar to CAR T cells, BiTE^®^ molecules trigger a polyclonal T cell response toward a specific antigen in an MHC independent manner^15,16^. FAP targeted BiTE^®^ molecules were generated by linking a single chain variable fragment (scFv) that recognizes and cross-ligates mouse CD3ε (K_d_ = 16-20 nM) on T cells to an scFv that recognizes mouse FAP (K_d_ = 0.1-0.3nM) (**Fig. 2A**).

**Figure 2.**
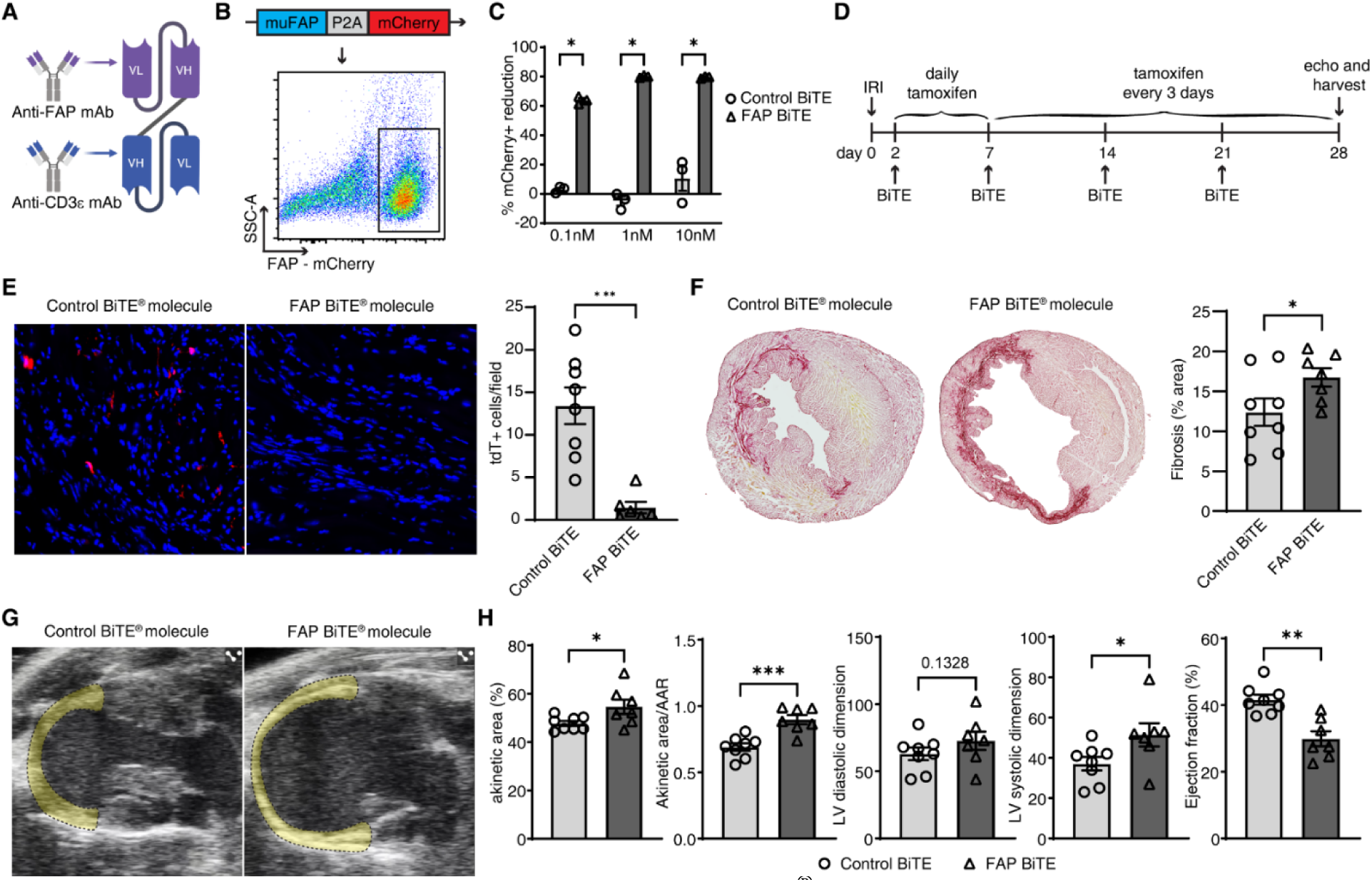
Anti-FAP bispecific T cell engagers (BiTE^®^ molecules) direct T cells to kill FAP^+^ cells but worsen cardiac function, exacerbate adverse chamber remodeling, and increase scar size after IRI. A) Structure of FAP BiTE^®^ molecule B) Generation of FAP^+^ mCherry reporter cell line for *in vitro* killing assays C) FAP BiTE^®^ molecule *in vitro* killing assay with naïve T cells D) Experimental schematic E) tdTomato expression and quantification in control vs. FAP BiTE^®^ molecule treated mice after IRI F) Picro Sirius Red staining and quantification G) Representative echocardiogram stills in systole with akinetic area highlighted H) Echocardiogram quantification

To test the ability of our FAP targeted BiTE^®^ molecule to eliminate FAP^+^ cells, we generated an immortalized target line by transducing k562 cells with a construct encoding *muFap* – P2A – mCherry, allowing for co-expression of mouse FAP and mCherry (**Fig. 2B**). Co-culture of naïve T cells with FAP-mCherry target cells resulted in robust elimination of FAP reporter cells only in the presence of FAP BiTE^®^ molecule, but not with the control BiTE^®^ molecule (isotype scFv, anti-CD3ε scFv), in a dose dependent manner (**Fig. 2C**).

To measure FAP^+^ cardiac fibroblasts depletion efficiency *in* vivo, we administered Fap tdT mice either control or FAP BiTE^®^ molecule following IRI (400µg/kg dosed weekly starting on day 2 post injury) (**Fig. 2D**). We chose this dose based on modeling of *in vivo* PK data to maintain a trough concentration >20X the *in vitro* determined EC_90_ (**fig. S2**)^17^. By measuring myocardial tdTomato immunofluorescence, we observed >95% target cell depletion (**Fig. 2E**). Despite robust target cell elimination, FAP BiTE^®^ molecule treated mice displayed larger infarct size and fibrosis on day 28 after IRI compared to those treated with the control BiTE^®^ molecule (**Fig. 2F**). Echocardiography revealed that FAP BiTE^®^ molecule treated animals demonstrated increased akinetic area, more extensive LV chamber dilation, and reduced ejection fraction (**Fig. 2G-H**). These results demonstrate a potential fallibility of T cell engaging therapies in the heart and suggest the existence of unintended effects that worsen cardiac function and remodeling to a degree that is sufficient to negate the beneficial effects of FAP*^+^* cell depletion. To this end, we sought to identify mechanisms that drive the deleterious effects of FAP BiTE^®^ molecules in the heart.

### FAP BiTE^®^ molecules reshape the cardiac T cell landscape

To investigate how FAP BiTE^®^ molecules lead to impaired cardiac function following IRI, we performed bulk RNA sequencing to ascertain tissue-level transcriptional differences in the hearts of FAP BiTE^®^ molecule versus control BiTE^®^ molecule treated mice after IRI. Bulk RNA-sequencing (day 8 following IRI) revealed numerous differentially expressed genes between treatment groups (**Fig. 3A**). Gene ontology (GO) enrichment analysis indicated that response to interferon gamma (IFNγ) was the most upregulated pathway in FAP BiTE^®^ molecule treated hearts compared to controls (**Fig. 3B**). RT-qPCR for *Ifng* and its downstream targets *Cxcl9* and *Ccl5*^18–20^ provided corroborating evidence that FAP BiTE^®^ molecules triggered activation of IFNγ signaling in the heart across multiple timepoints (**Fig. 3C**). This response appeared to be restricted to the heart, as we did not observe systemic IFNγ in the periphery (**fig. S3**).

**Figure 3.**
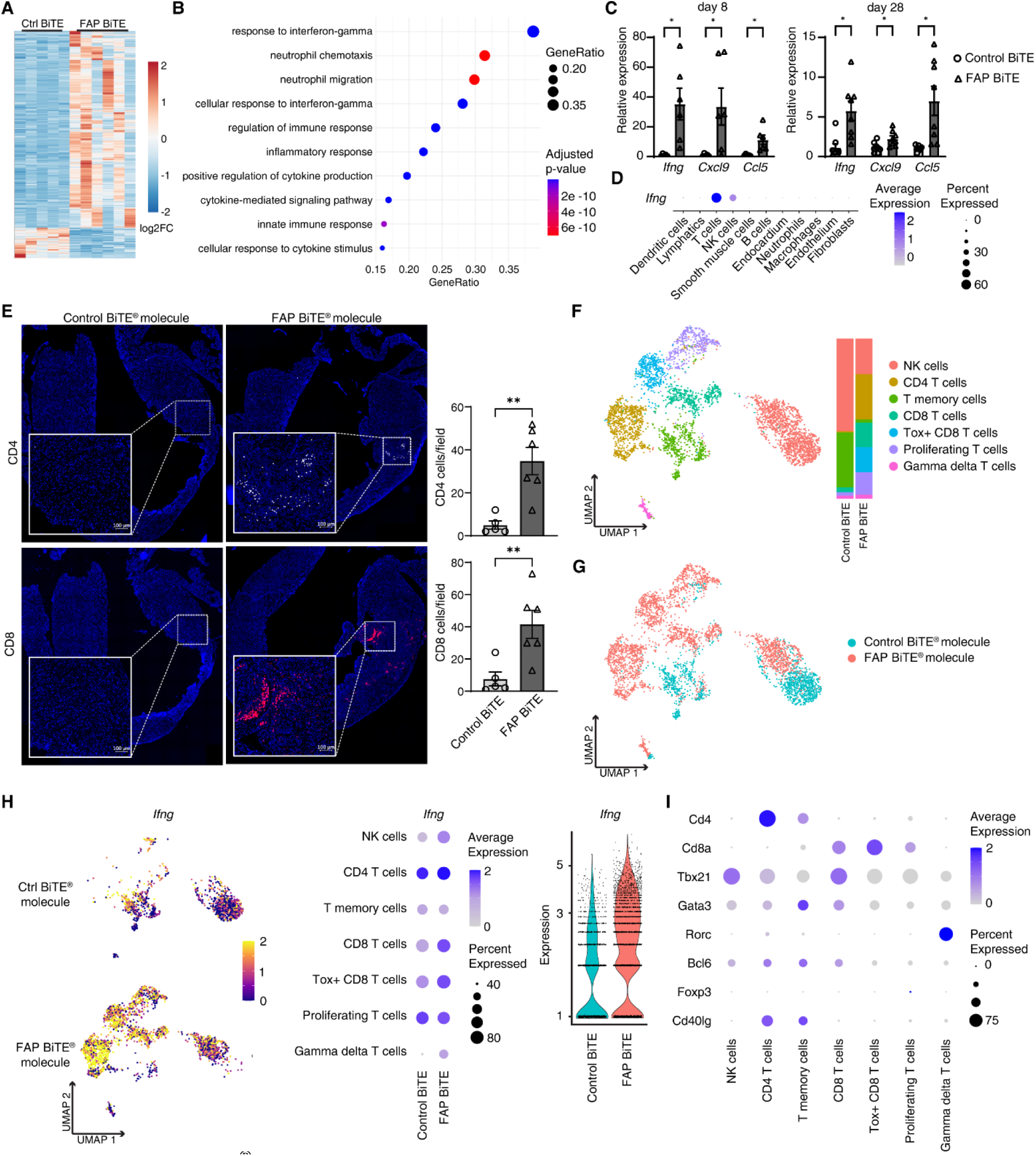
FAP BiTE^®^ molecule treatment increases the frequency of cardiac T cells and induces distinct T cell states. A) Bulk RNA-sequencing heat map of differentially expressed genes between control and FAP BiTE^®^ molecule treated hearts B) GO pathway analysis of FAP BiTE^®^ molecule treated hearts compared to control in bulk RNA-sequencing C) RT-qPCR of *Ifng, Cxcl9, and Ccl5* between conditions at day 8 and day 28 after IRI D) Dot plot of *Ifng* expression between cell types in single cell RNA sequencing of BiTE^®^ molecule treated hearts day 8 after IRI E) CD4 and CD8 T cell staining and quantification in control vs FAP BiTE^®^ molecule treated mice after IRI F) Global UMAP of T and NK cells and stack plot split by condition in BiTE^®^ molecule treated hearts day 8 after IRI G) Feature plot of cells split by condition H) Feature plot and dot plot of *Ifng* split by condition and subcluster. Violin plot of *Ifng* expression split by condition I) Dot plot of T cell surface marker and transcription factors split by cluster

To further define the source and recipients of IFNg signaling and resultant effects on the cardiac landscape, we isolated live cells by sorting DAPI^-^ DRAQ5^+^ cells from enzymatically digested hearts from mice treated with either control or FAP BiTE^®^ molecules on day 8 following IRI and constructed single cell RNA sequencing libraries using the 10X Genomics platform (**fig. S4A**). Single-cell libraries were sequenced, aligned to the mouse reference genome, and underwent QC filtering (**fig. S4B-D**). We then performed unsupervised clustering and differential expression analysis using Seurat (**fig. S5A**). We identified 11 clusters, each representing a unique cell type (**fig. S5B-C**). UMAP representation revealed obvious differences between experimental conditions across cell types (**fig. S5D**). Interrogation of *Ifng* mRNA expression indicated that T cells represented the primary source of this cytokine in FAP BiTE^®^ molecule treated hearts (**Fig 3D**). This finding is suggestive of T cell activation within the myocardium ^21^.

We next performed immunofluorescence staining for CD4 and CD8 in control and FAP BiTE^®^ molecule treated hearts on day 8 after IRI. We observed a robust increase in the number of both CD4^+^ and CD8^+^ T cells, which were most prominent in the region of the infarct closest to the border zone (**Fig. 3E**). Consistent with increased T cell abundance and activation, T cell activation markers and downstream T cell receptor signaling components were amongst the highest differentially expressed transcripts identified by bulk RNA sequencing of hearts from FAP BiTE^®^ molecule treated mice (**fig. S6**). To further characterize how FAP BiTE^®^ molecules influence T cell phenotypes, we performed additional single cell RNA sequencing of T/NK cell enriched preparations (CD45^+^ CD64^-^ Ly6G^-^ CD19^-^ cells) from control and FAP BiTE^®^ molecule treated hearts on day 8 following IRI (**fig. S7A-D**). Unsupervised clustering and differential expression analysis using Seurat identified 7 clusters (**Fig. 3F**), each representing a unique T cell state (**fig. S8A-C**). UMAP representation revealed major shifts between experimental conditions across T and NK cell states (**Fig. 3G**). Namely, control hearts contained predominately NK cells and T memory cells, a cluster marked by expression of *Tcf7*, *Ccr7*, and *Il7r*. In the FAP BiTE^®^ molecule condition, we observed expansion of CD4 T cells, CD8 T cells, *Tox^+^* CD8 T cells, and proliferating T cells. NK cells were reduced in relative abundance and T memory cells were almost entirely absent. Pathway analysis of differentially expressed genes between conditions revealed that the top enriched pathways were predominantly associated with cell cycle regulation (**Fig. S8D**).

We analysed *Ifng* expression between T/NK cell states and conditions. This dataset also demonstrated higher *Ifng* expression in FAP BiTE^®^ molecule compared to control T/NK cells. Within the T/NK cell compartment of FAP BiTE^®^ molecule treated hearts, *Ifng* was most enriched in CD4^+^ T cells and to a lesser extent in CD8^+^ T cells including *Tox^+^* CD8^+^ T cells (**Fig. 3H**). We also observed that the expanded population of CD4^+^ T cells found within FAP BiTE^®^ molecule treated hearts had a greater proportion of *Cd40lg* expressing cells than the CD4^+^ cluster in the control BiTE^®^ molecule group (**Fig. 3I).** It is well described that CD40L signaling to CD40 on macrophages and other antigen presenting cells can augment T cell activation and production of IFNγ either through inducing macrophage production of IL-12^22,23^, a T_H_1 specifying cytokine^24^, or by inducing expression of costimulatory molecules CD80/86^25,26^. Consistent with this concept, we have shown that CD40 agonists can instruct cardiac macrophages to recruit T cells and elicit IFNγ production^27^. This offers a potential mechanism by which T cell-centered IFNγ signaling can be amplified or sustained following FAP BiTE^®^ molecule treatment. Indeed, IFNγ and CD40L-CD40 signaling have been shown to be drivers of adverse cardiac outcomes in models of myocarditis^27,28^.

### Blockade of IFN-γ and CD40L unmasks therapeutic efficacy of FAP BiTE^®^ molecules

We next set out to delineate whether IFNγ and CD40L signaling are responsible for the adverse cardiac effects observed following FAP BiTE^®^ molecule treatment. As conflicting evidence exists pertaining to whether IFNγ^29–33^ and CD40L^34,35,36^ signaling are necessary for T cell mediated target killing, we investigated the requirement for these pathways in FAP BiTE^®^ molecule mediated target cell elimination. We confirmed production of IFNγ and expression of CD40L by T cells in response to FAP BiTE^®^ molecule in our *in vitro* killing assay (**fig. S9A-B**). Neutralization of IFNγ or CD40L did not compromise FAP BiTE^®^ molecule target killing *in vitro* (**fig. S9C**). Instead, we observed that blocking TRAIL but not FasL reduced the ability of the FAP BiTE^®^ molecule to kill target cells (**fig. S9D**). These data supported the possibility that blocking IFNγ and CD40L *in vivo* could attenuate harmful collateral inflammation induced by FAP BiTE^®^ molecules without impacting target cell killing and potentially unmask a therapeutic benefit.

To test this hypothesis, we treated Fap tdT reporter mice with either control or FAP BiTE^®^ molecules in the presence of isotype control or anti-IFNγ + anti-CD40L antibodies after IRI (**Fig. 4A**). To assess the impact of these interventions on IFNγ signaling, we measured *Ifng* and *Cxcl9* expression by qRT-PCR. We observed increased in *Ifng* and *Cxcl9* expression in FAP BiTE^®^ molecule compared to control BiTE^®^ molecule treated hearts, which were both reduced by anti-IFNγ + anti-CD40L antibodies (**Fig. 4B**). Next, we performed immunofluorescence for tdTomato expression to assess *in vivo* target cell elimination. Importantly, anti-IFN-γ + anti-CD40L antibody treatment did not interfere with FAP BiTE^®^ molecule mediated target cell killing (**Fig. 4C**) consistent with our *in vitro* assays. Measurement of infarct fibrosis 28 days after IRI revealed that neutralization of IFNγ and CD40L signaling led to reductions in fibrosis compared to isotype treated mice administered FAP BiTE^®^ molecules (**Fig. 4D**). Echocardiography demonstrated that neutralization of IFNγ and CD40L signaling similarly reduced akinetic area, attenuated LV chamber dilation, and improved ejection fraction in FAP BiTE^®^ molecule treated mice (**Fig. 4E-F**). IFNγ and CD40L neutralization had no observed effect on fibrosis, akinetic area, LV dimensions, or ejection fraction in mice that received the control BiTE^®^ molecule. Strikingly, IFNγ and CD40L blockade unmasked a therapeutic benefit of FAP BiTE^®^ molecules, as evidenced by reductions in akinetic area and LV chamber dimensions and increased ejection fraction when compared to mice that received the control BiTE^®^ molecule. Evaluation of the individual contribution of IFNγ and CD40L signaling blockade on cardiac remodeling in FAP BiTE^®^ molecule treated mice revealed that anti-IFNγ and anti-CD40L antibodies did not impact FAP cell killing. While both IFNγ and CD40L neutralization led to improvements in cardiac fibrosis and cardiac remodeling, IFNγ inhibition had a greater effect (**fig. S10**). Collectively, these data indicate that collateral activation of IFNγ and CD40L pathways contributes to the detrimental effects of FAP BiTE^®^ molecules.

**Figure 4.**
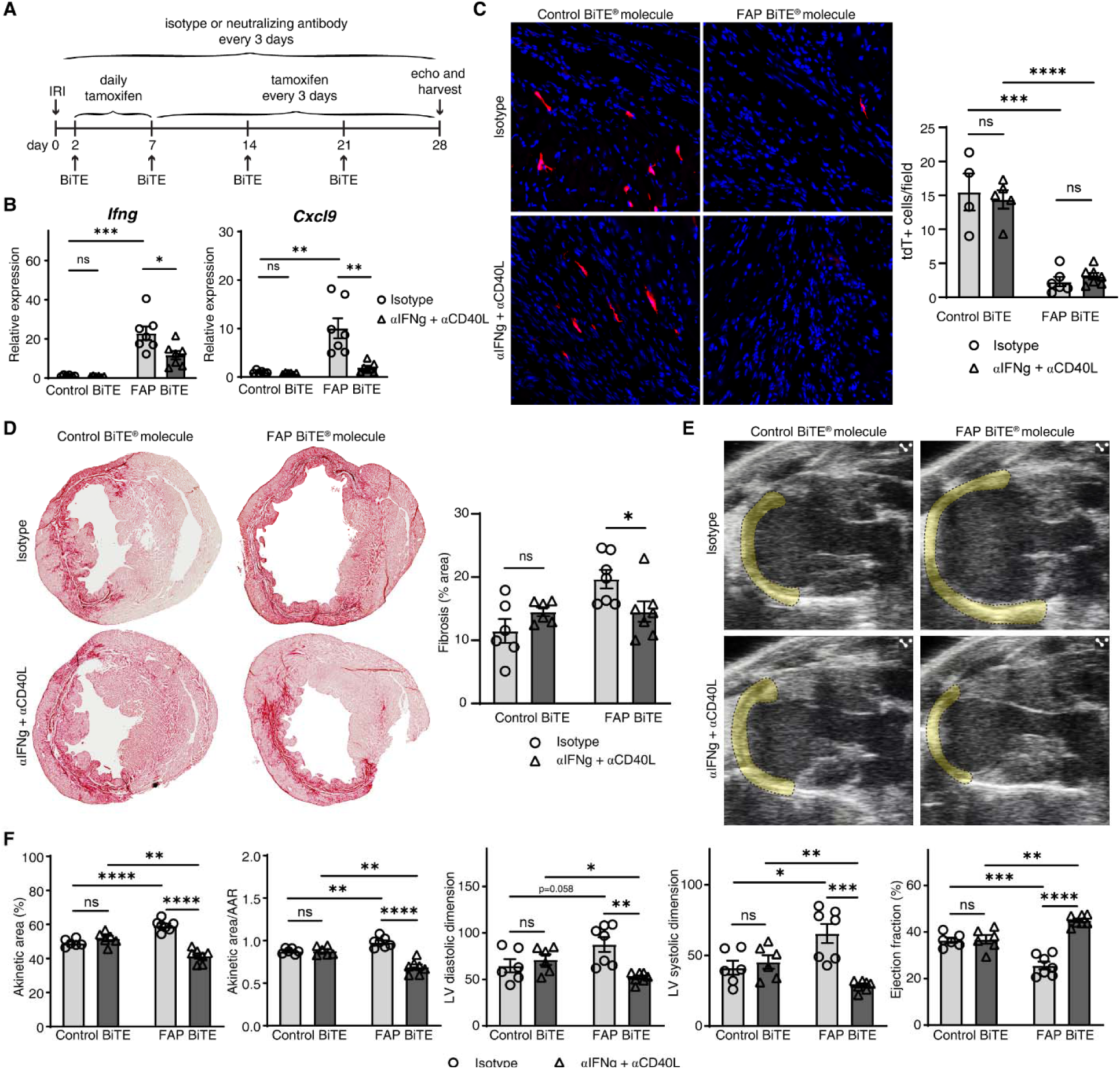
IFNg and CD40L neutralization preserves target killing with FAP BiTE^®^ molecule while preventing adverse effects after cardiac IRI. A) Experimental schematic B) RT-qPCR of *Ifng* and *Cxcl9* between conditions at day 28 after IRI C) tdTomato expression and quantification in FAP BiTE^®^ molecule vs control BiTE^®^ molecule treated mice after IRI with and without IFNg + CD40L neutralization D) Picro Sirius Red staining and quantification E) Representative echocardiogram stills in systole with akinetic area highlighted F) Echocardiogram quantification

### FAP BiTE^®^ molecules drive the emergence of an alternative lineage of activated fibroblasts

To understand how FAP BiTE^®^ molecule-mediated T cell activation accelerates adverse cardiac remodeling and increases fibrosis, we further analysed our single cell RNA sequencing dataset. Pathway analysis of differentially expressed genes between conditions identified IFNγ signaling as the most upregulated pathway across major cell types in FAP BiTE^®^ molecule treated mice compared to controls (**fig. S11A**). Differential expression analysis revealed that the greatest number of genes differentially expressed between experimental conditions resided in fibroblasts followed by endothelium, macrophage, and neutrophil populations (**fig. S11B**). To discern which cell types are activated by IFNγ, we focused our analysis on *Cxcl9* expression, a specific marker of IFNγ signaling^37,38^. While many cell types displayed an increase in the relative expression level and proportion of cells expressing *Cxcl9* in the FAP BiTE^®^ molecule group compared to controls, fibroblasts, macrophages, and dendritic cells demonstrated the greatest enrichment (**fig. S11C-D**). To ascertain the spatial context of IFNg activation within the heart after FAP BiTE^®^ molecule treatment, we performed Xenium spatial transcriptomics on sections obtained from mice day 8 after IRI (**fig. S11E**). Cell annotation and analysis of spatial transcriptomic data indicated that cell types expressing Cxcl9 were located within the infarct (**fig. S11F-I**).

We then performed detailed subclustering of fibroblasts and macrophages, as they represented abundant cell populations that expressed the largest number of differentially expressed genes between conditions. We identified 8 distinct transcriptional states of cardiac fibroblasts that markedly varied across control and FAP BiTE^®^ molecule conditions (**Fig. 5A-C, S12A**). As anticipated FAP BiTE^®^ molecule treatment resulted in elimination of FAP fibroblasts (cluster 1). Pathway analysis of top differentially expressed genes between conditions revealed complement, oncostatin M, IFNγ, and IL-1 regulation of extracellular matrix as the most enriched pathways in FAP BiTE^®^ molecule treated fibroblasts (**Fig. S12B**). Feature plots of *Cxcl9* expression split by condition revealed upregulation of *Cxcl9* in the FAP BiTE^®^ molecule group within fibroblast clusters 4 and 7, which were uniquely represented in the FAP BiTE^®^ molecule group (**Fig. 5D-E**). Fibroblast clusters 4 and 7 did not express *Fap* at levels that could be detected by single cell RNA sequencing. *Fap* expression was reduced in the FAP BiTE^®^ molecule group compared to controls (**Fig. 5F-G**). Intriguingly, fibroblast clusters 4 and 7 expressed *Postn* indicating that they represent an activated fibroblast cell state (**Fig. 5H-I**). RNA velocity trajectory analysis revealed vector convergence within cardiac fibroblast clusters 1, 4, and 7 suggesting that they represent the most terminal states (**Fig. 5J**). NicheNet ligand-receptor analysis^39^ focused on signaling to fibroblasts predicted T cell derived IFNγ as the predominate incoming signal to fibroblasts in the heart following FAP BiTE^®^ molecule treatment (**Fig 5K**). We leveraged our Xenium spatial transcriptomic data to illuminate the spatial location of activated fibroblasts in control and FAP BiTE^®^ molecule treated hearts after IRI. *Fap^+^* fibroblasts were present within the infarct of control mice and eliminated from FAP BiTE^®^ molecule treated mice. IFNγ activated fibroblasts were uniquely present within the infarct of FAP BiTE^®^ molecule treated hearts (**Fig. 5L, S12C**). Collectively, these findings uncover the emergence of a FAP^-^ activated fibroblast state within the infarct of FAP BiTE^®^ molecule treated animals that are predicted to arise from T cell – fibroblast IFNγ signaling.

**Figure 5.**
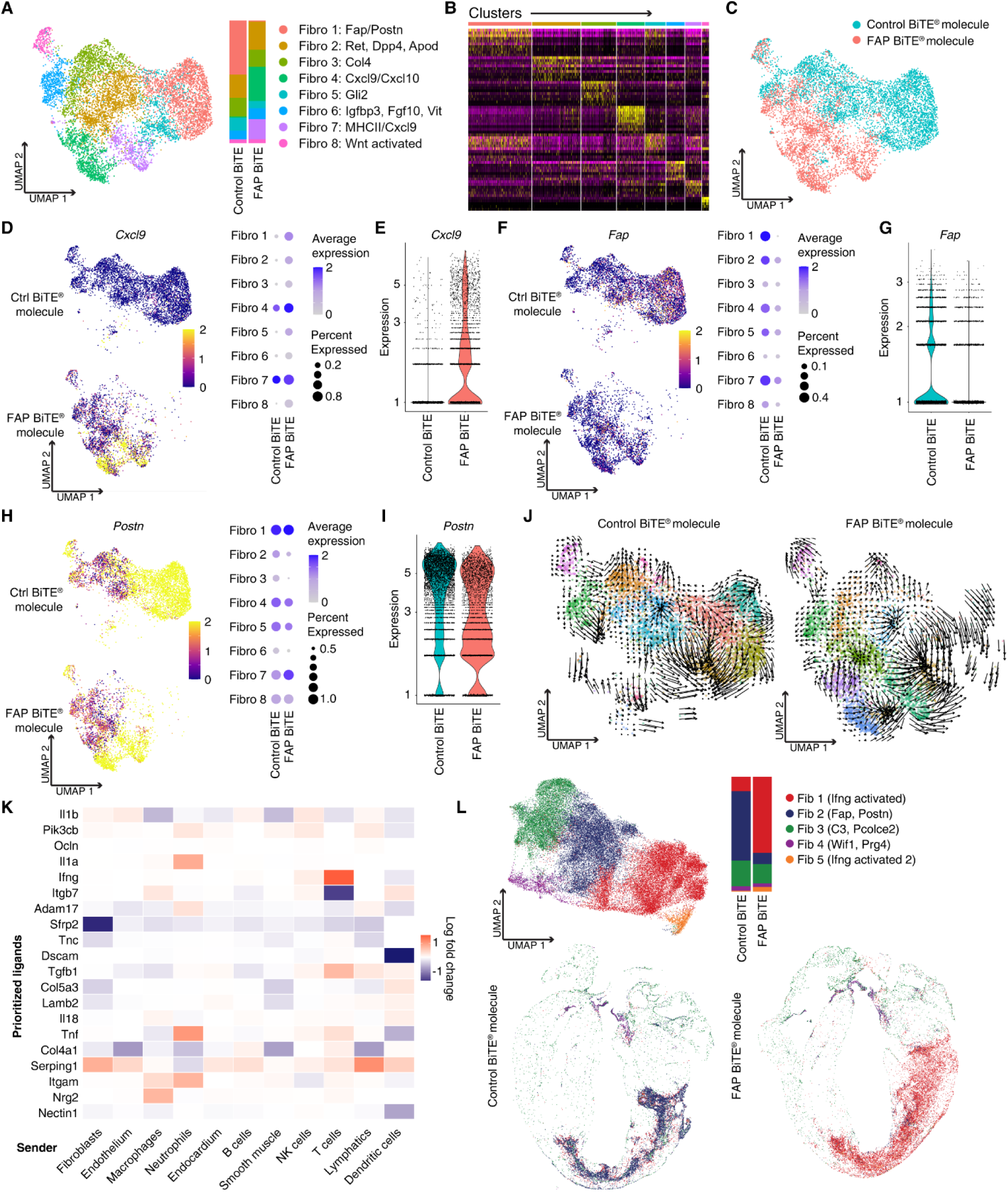
Fibroblast subclustering reveals reduction of target cells with expansion of IFNg responsive fibroblasts in FAP BiTE^®^ molecule treated hearts. A) UMAP of subclustered fibroblasts and stack plot split by condition in BiTE^®^ molecule treated hearts day 8 after IRI B) Heatmap of differentially expressed genes by cluster C) Feature plot of cells split by condition D) Feature plot and dot plot of *Cxcl9* split by condition and subcluster E) Violin plot of *Cxcl9* F) Feature plot and dot plot of *Fap* split by condition and subcluster G) Violin plot of *Fap* H) Feature plot and stack plot of *Postn* split by condition and subcluster I) Violin plot of *Postn* J) RNAvelocity of fibroblast subclustering UMAP split by condition K) NicheNet receptor ligand analysis heatmap with fibroblast as receiver L) Xenium spatial transcriptomic UMAP of fibroblast subclustering with stack plot split by condition

Macrophage subclustering identified 7 distinct transcriptional cell states (**fig. S13A-C**) that markedly shift between FAP BiTE^®^ molecule and control groups (**fig. S13D**). Pathway analysis revealed matrix metalloproteinases, IFNγ signaling, IL4 regulation of apoptosis, leptin, and thymic stromal lymphopoietin as the most enriched pathways in FAP BiTE^®^ molecule treated macrophages compared to control (**fig. S13E**). *Cxcl9* expression was increased in FAP BiTE^®^ molecule treated macrophages and enriched in cluster 4 (**fig. S13F-G)**. NicheNet ligand-receptor analysis of incoming signals to macrophages identified T cell-derived IFNγ and CCL3 as major modes of T cell – macrophage communication following FAP BiTE^®^ molecule treatment (**fig. S13H**). Interestingly, cluster 4, the *Cxcl9* expressing macrophages that are expanded in the FAP BiTE^®^ molecule condition, also expresses the highest levels of CD40 (**fig. S13I**). Xenium spatial transcriptomic analysis of immune cells in FAP BiTE^®^ molecule treated hearts demonstrated the expansion of IFNγ activated macrophages throughout the infarct (**fig. S13J-K**).

### IFNγ signaling to fibroblasts drives adverse remodeling and fibroblast activation following FAP BiTE^®^ molecule treatment

The above single cell RNA sequencing and spatial transcriptomics data indicated emergence of a trajectory of activated *Postn^+^* fibroblasts potentially driven by T cell derived IFNγ signaling. Based on these findings, we next tested the hypothesis that blocking IFNγ signaling to fibroblasts may be sufficient to mitigate the adverse effects of FAP BiTE^®^ molecules. We generated control and *Pdgfra^ert2Cre^Ifngr1^flox/flox^*mice and treated them with FAP BiTE^®^ molecules following IRI. Tamoxifen was administrated to all mice beginning after IRI (**Fig. 6A**). We observed a reduction in fibrosis in the knockout animals (**Fig. 6B).** Deletion of the IFNγ receptor from fibroblasts was also sufficient to reduce akinetic, decrease LV volumes, and increase ejection fraction (**Fig. 6C-D**). Together, these data demonstrate that IFNγ signaling to fibroblasts constitutes a critical mechanism by which FAP BiTE^®^ molecules contribute to accelerated cardiac remodeling, enhanced fibrosis, and deterioration of LV function.

**Figure 6.**
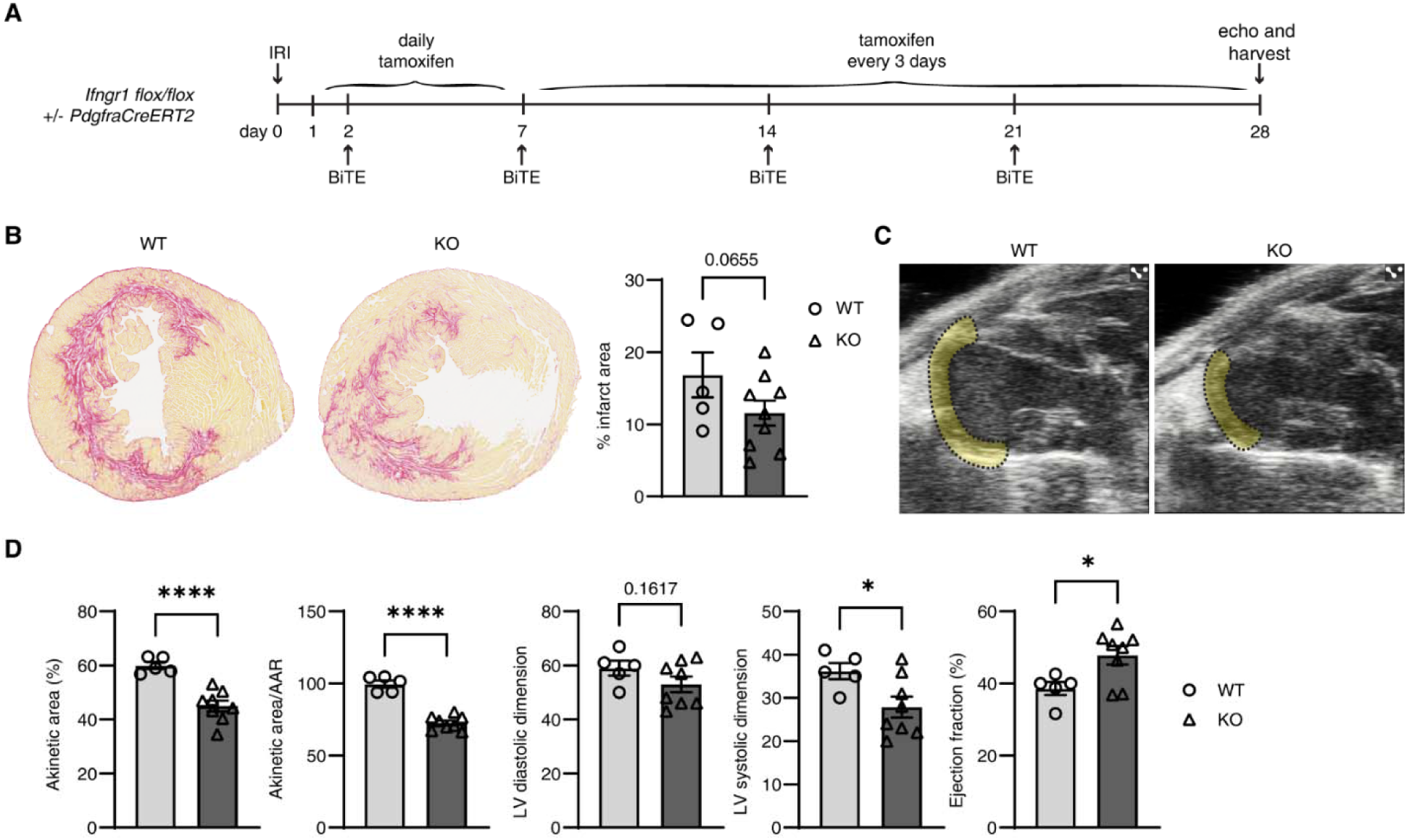
Genetic elimination of IFNg signaling in fibroblasts rescues adverse cardiac phenotype in FAP BiTE^®^ molecule treated hearts after IRI. A) Experimental schematic B) Picro Sirius Red staining and quantification C) Representative echocardiogram stills in systole with akinetic area highlighted D) Echocardiogram quantification

## Discussion

Previous work has established that FAP^+^ fibroblasts represented an activated cell state associated with cardiac fibrosis and adverse clinical outcomes^40,41^. We build on this foundation and show that FAP^+^ fibroblasts are specified in the mouse heart following reperfused MI and that depleting these cells and their progeny results in reduced scar size and improved cardiac function, highlighting the therapeutic potential of depleting FAP^+^ fibroblasts from the heart. Recent technological advancements have now made it possible to remove specific cell populations in humans through engineered cell products and antibody-based therapies.

T cell engaging therapies including CAR T cells and BiTE^®^ molecules are now commonly used in oncology settings to remove cancerous cells and have the potential to reshape treatment of rheumatologic conditions by depleting auto-reactive immune cells^42^. Here, we show that FAP BiTE^®^ molecules enable robust and specific depletion of FAP^+^ fibroblasts from the injured heart. Unexpectedly, they also induce collateral myocardial inflammation by activating T cells and macrophages through an IFN-γ and CD40 dependent mechanism, which results in specification of an activated fibroblast lineage that is distinct from FAP^+^ fibroblasts or myofibroblasts. We further demonstrate that blockade of IFN-γ and CD40 signaling attenuates the harmful immunomodulatory effects of the FAP BiTE^®^ molecules without impacting their ability to deplete FAP^+^ fibroblasts. As a consequence, inhibition of IFN-γ and CD40 signaling unmasked the therapeutic benefit of FAP BiTE^®^ molecules leading to improved LV systolic function and reduced post-MI remodeling. Finally, we demonstrate that IFN-γ signaling to fibroblasts is a central mechanism by which FAP BiTE^®^ molecules augment cardiac pathology.

Our study is not without limitations. The FAP BiTE^®^ molecule used in this work displays high affinity for both FAP and CD3 and may result in greater T cell activation compared to other constructs. In addition, we dosed the anti-FAP BiTE^®^ molecule to achieve plasma trough levels >20X of the EC_90_ to ensure adequate cardiac target coverage. It is possible that lower doses may lead to less IFN-γ and CD40 signaling. Finally, our FAP BiTE^®^ molecule has the capacity to engage a wide range of CD4 and CD8 T cells. Improved understanding of the T cell subsets that reside within target tissues and are engaged by BiTE^®^ molecules will likely be instrumental to optimize this therapeutic approach. While diverse subsets of T cells occupy the ischemic and failing heart, CD4 T cells with T_h_1 properties are abundant and known to contribute to adverse cardiac remodeling^43–46^. Engagement of similar CD4 T cells with limited cytotoxic activity likely triggered IFN-γ and CD40 signaling and accelerated disease pathogenesis, which may be avoided by more selective BiTE^®^ molecules that engage CD8 T cells or other CD4 T cell subsets with less pronounced inflammatory potential.

In conclusion, our findings elucidate a previously unrecognized liability of BiTE^®^ molecules in the heart, identify mechanisms responsible for collateral inflammation, and will ultimately pave the way for new strategies that maximize efficacy and optimize patient safety of T cell-engaging immunotherapies for cardiovascular disease.

## Methods

### Animal models

Animal studies were performed in compliance with guidelines set forth by the National Institutes of Health Office of Laboratory Animal Welfare and approved by the Washington University institutional animal care and use committee. Animals were housed in a controlled environment with a 12 h light–dark cycle, with free access to water and a standard chow diet. Strains used were *Rosa26^Lox-STOP-Lox-DTR^ (*JAX # 007900), *Rosa26^Lox-STOP-Lox-DTR^* (JAX #007914), and *Fap^ert2Cre^*(JAX #040069) on the C57BL/6 background. Experiments were performed on mice 2–5 months of age and individual experiments contained mice of similar ages in all experimental conditions. Similar numbers of male and female mice were used for experiments. For Cre recombination, mice were orally gavaged with 120mg/kg of tamoxifen (Sigma #T5648) at the indicated frequency. For DTR depletion, mice were injected IP with 200 ng DT (Sigma #D0564). For BiTE^®^ molecule experiments, mice were injected IP with 400ug/kg of either control or FAP BiTE^®^ molecule. For the IFNg and CD40L neutralization experiments, mice were injected IP with the 300ug designated antibody they were randomized to. Antibodies used are all from Bioxcell: IgG1 isotype control (#BE0088), IFNg (#BE0054), CD40L (#BE0017-1).

### Ischemia reperfusion injury

Echo guided ischemia reperfusion was performed as previously described. In brief, mice were anesthetized, intubated and mechanically ventilated. 2 micromanipulator-mounted hypodermic 30G needles were inserted through the left side chest wall and into the myocardium ventral and dorsal to the left anterior descending artery. Both needles were pinched together to induce ischemia for 60 minutes before being released to induce reperfusion. During this period of ischemia, the area at risk (AAR) is measured by quantifying the akinetic area during the 60 minutes of ischemia.

### Echocardiography

Mice were sedated with Avertin (0.005 ml g−1), and two-dimensional and motion mode images were obtained in the long and short axis views 4 weeks after MI using the VisualSonics770 Echocardiography System. Left diastolic volume, left systolic volume, akinetic area and total LV area were measured using edge detection and tracking software (VivoLab). The EF was calculated as (left diastolic volume − left systolic volume)/left diastolic volume. The akinetic percentage was calculated as the akinetic area/total LV area.

### Immunofluorescence and immunohistochemistry

Mice were euthanized, and the hearts perfused with PBS. For chronic timepoint, day 28 post-IRI experiments, hearts were cut into thirds along the transverse plane and fixed overnight with 4% PFA in PBS at 4C. They were then dehydrated in 30% sucrose in PBS overnight at 4C. The hearts were embedded in optimal cutting temperature compound (Sakura, 4583) and frozen at -80 °C for 30 min. Hearts were then sectioned at 7um using a Leica cryostat, mounted on positively charged slides, and stored at -80C. Slides were brought to room temperature (RT) for 5 min and washed in PBS for 5 min. For tdTomato imaging, slides were immediately mounted with mounting media with DAPI (Invitrogen #P36931). For CD4 (Abcam #ab183685) and CD8 (Abcam #ab217344) staining, slides were fixed in 10% methanol with 3% H2O2 in PBS for 20 minutes at RT before washing in PBS 3 x 3 minutes. Sections were then blocked in 10% BSA in PBS at for 30 minutes at RT. Primary antibodies were diluted 1:1000 (CD4) or 1:200 (CD8) in 1% BSA PBS and added to sections for 1 hour at RT. Slides wee washed 3 x 3 minutes before the anti-Rb-HRP secondary antibody (Akoya #ARR1001KT) was added for 10 minutes at RT. Slides were then washed in PBS 3 x 3minutes before Opal 520 (Akoya FP1487001KT) was diluted 1:200 and added to sections for 10 minutes at RT. Slides were washed in PBS 3 x 3 minutes before mounting slides with DAPI. For picrosirius red staining, the PSR stain kit (Cardiac Muscle (Abcam #ab245887) kit was used before mounting with Cytoseal 60 (Mercedes #FIS 83104). Images were taken on the ZEISS Axioscan 7 slide scanner. Quantification of cell numbers was performed in Zeiss Zen, and data are displayed as the average of each 20× field per mouse.

### BiTE^®^ molecule generation

BiTE^®^ molecules used in this study include: (A) 64281 muFAPa BiTE^®^ molecule containing a mouse anti-mouse FAPa scFv derived from the 73.3_MABC1145 antibody; and (B) 64280 control BiTE^®^ molecule containing an anti-idiotype scFv that does not recognize a known target. Both BiTE^®^ molecules also contain the KE3 BiTE^®^ molecule consisting of a mouse anti-mouse CD3e KE3 scFv derived from the KT3 mAb sequence. BiTE^®^ molecule HLE molecules were produced using standard protocols described previously1. Briefly, BiTE^®^ molecule HLE molecules were expressed from CHO stable cell lines using serum-free media and purified from cell culture supernatant using protein A capture (Mabselect Sure, Cytiva) followed by preparative size-exclusion chromatography (Superdex 200, Cytiva). Final product quality was confirmed by mass spectrometry (ThermoFisher Vanquish LC / Q Exactive Plus Orbitrap), analytical size exclusion chromatography (HPLC-SEC, Agilent 1100 Series HPLC) and endotoxin testing (EndoSafe^®^-MCS, Charles River Laboratories).

### BiTE^®^ molecule pharmacokinetics analyses

Analysis of serum samples was performed using electro-chemiluminescent immunoassays on the MSD Sector 600 instrument (Meso Scale Diagnostics, Rockville, MD). Biotinylated recombinant mouse FAP protein (R&D Systems, Cat# 8647-SE) was coated onto Streptavidin SECTOR plates as the capture reagent. The standards and samples were diluted in assay buffer (Blocker™ BLOTTO in TBS) and then added to the plate and incubated for one hour, followed by washing and then incubation with a ruthenylated anti-mouse IgG, Fcγ subclass 1 specific antibody (JacksonImmuno Research, cat# 115-005-205) as the detection reagent. The analyte serum concentrations were interpolated from standard curves using the corresponding analyte prepared in pooled mouse serum. Noncompartmental analyses to obtain PK parameters were calculated via WatsonLIMS software (Thermo Fisher) using the mean concentration of the grouped mice at each time point.

### *In vitro* killing assay

FAP–mCherry lentiviral particles were generated using a third-generation lentiviral vector backbone. The viral supernatant was collected 48 hours post-transfection and concentrated by overnight centrifugation at 8,000 rpm at 4 °C. The pellet was resuspended in RPMI-1640 medium. 2e6 K562 (ATCC #CCL243) cells were transduced with 200 µL of the viral supernatant and maintained in culture for 7 days. mCherry+ cells were subsequently enriched by fluorescence-activated cell sorting (FACS) in the institutional flow cytometry core. mCherry+ target cells were plated at a density of 400,000/mL in a 96well round bottom plate (Corning #3879) with RPMI 1640 medium (Gibco #11875085) supplemented with 10% fetal bovine serum (Corning #35-010-CV) and 100U/mL of Penicillin-Streptomycin (Gibco #15140122). T cells were isolated from spleens using a pan T cell isolation kit (Miltenyi #130-095-130) and plated with the target cells at 4million cells/mL for an effector to target ratio of 10:1. The desired concentration of control or FAP BiTE^®^ molecule was added and the killing assay was allowed to proceed for 48hours before flow cytometric analysis on the Cytek Aurora. Killing was normalized to the number of mCherry positive cells in wells containing the target cells and T cells but no BiTE^®^ molecule. Conditions were plated in triplicate. Media was collected at time of analysis for IFNg ELISA (R&D #DY485).

### RT-qPCR

RNA was isolated from snap frozen heart tissue by lysing tissue in 1mL of TRIzol reagent (Invitrogen #15596026) with one 5mm stainless steel bead (Qiagen #69989) in the TissueLyser LT (Qiagen #85600) for 10 minutes at 50Hz. 200uL of chloroform (Sigma #151823) was added to the homogenate, the mixture was vortexed and left at RT for 5 minutes before centrifuging at 12000rcf for 15 minutes. The aqueous phase was isolated and purified using the PureLink RNA Mini Kit (Invitrogen #121830181A). cDNA was generated using the High-Capacity cDNA Reverse Transcription Kit (Fisher #43-688-14). qPCR reactions were prepared with PowerUP SYBR Green Master Mix (Thermo #A25742). Primer sequences were as follows: *Ccl5* fwd: GGGTACCATGAAGATCTCTGC rev: TCTAGGGAGAGGTAGGCAAAG *Cxcl9* fwd: GTTCGAACCCTAGTGATAAG rev: GTTTGAGGTCTTTGAGGGATTTG *Ifng* fwd: GGCCATCAGCAACAACATAAG rev: GTTGACCTCAAACTTGGCAATAC *36b4* fwd: ATCCCTGACGCACCGCCGTGA rev: TGCATCTGCTTGGAGCCCACGTT Primers were purchased from Integrated DNA Technologies. qPCR was performed using QuantStudio 6 (Thermo Fisher Scientific). Expression was normalized to *36b4*.

### Bulk RNA sequencing

Purified RNA was isolated as above. Total RNA integrity was determined using Agilent Bioanalyzer or 4200 Tapestation. Library preparation was performed with 5 to 10ug of total RNA with a Bioanalyzer RIN score greater than 8.0. Ribosomal RNA was removed by poly-A selection using Oligo-dT beads (mRNA Direct kit, Life Technologies). mRNA was then fragmented in reverse transcriptase buffer and heating to 94 degrees for 8 minutes. mRNA was reverse transcribed to yield cDNA using SuperScript III RT enzyme (Life Technologies, per manufacturer’s instructions) and random hexamers. A second strand reaction was performed to yield ds-cDNA. cDNA was blunt ended, had an A base added to the 3’ ends, and then had Illumina sequencing adapters ligated to the ends. Ligated fragments were then amplified for 12-15 cycles using primers incorporating unique dual index tags. Fragments were sequenced on an Illumina NovaSeq X Plus using paired end reads extending 150 bases.Basecalls and demultiplexing were performed with Illumina’s bcl2fastq software with a maximum of one mismatch in the indexing read. RNA-seq reads were then aligned to the Ensembl release 101 primary assembly with STAR version 2.7.9a1. Gene counts were derived from the number of uniquely aligned unambiguous reads by Subread:featureCount version 2.0.32. Isoform expression of known Ensembl transcripts were quantified with Salmon version 1.5.23. Sequencing performance was assessed for the total number of aligned reads, total number of uniquely aligned reads, and features detected. The ribosomal fraction, known junction saturation, and read distribution over known gene models were quantified with RSeQC version 4.04.

### Single cell RNA-sequencing analysis

All live cells (experiment 1) or T and NK cells (experiment 2) from FAP BiTE^®^ molecule and control BiTE^®^ molecule mice were sorted 8 days after IRI. scRNAseq libraries were generated using the 10XGenomics 3′v4 platform. Libraries were sequenced using a NovaSeq at the McDonnel Genome Institute. Sequencing reads were aligned to the GRcm39 transcriptome using CellRanger (v9.0.1) from 10X Genomics. Downstream data processing was performed in the R package Seurat (v4). RNA count data was scaled and normalized using SCtransform. Quality control filters of genes per cell (>200, <8000), read counts per cell (<50000), and proportion of mitochondrial reads per cell (<10%), were applied (Sup. fig. 4 and 7). Principal component analysis was used for dimensionality reduction. Data were clustered at multiple resolutions and visualized using Uniform Manifold Approximation and Projection (UMAP) projections. Differential gene expression of upregulated genes across conditions and clusters was generated in Seurat using FindAllMarkers with (only.pos = TRUE, min.pct = 0.1, logfc.threshold = 0.25). Statistically significant genes (adj. p val < 0.05) from FindAllMarkers were used as input for pathway analysis in EnrichR. For RNA velocity analysis, CellRanger aligned BAM files were generated and utilized as an input for Velocyto CLI to obtain loom files with estimations of spliced and unspliced RNA counts. The loom files were merged with the annotated fibroblast subcluster object and pre-processed prior to estimation of RNA velocity via stochastic modelling. RNA velocity graphs were generated using velocyto.R.

### Xenium spatial transcriptomics

Raw sequencing data was aligned, with counts decoded into spatial transcripts and assigned to segmented cells. Cell segmentation was assigned via a neural network algorithm and a combination of nuclear, cell interior, and cell membrane stains. The resulting cell by gene matrix contains corresponding spatial coordinates for each cell, and was used for downstream analysis including clustering, differential expression analysis, and Uniform Manifold Approximation and Projection (UMAP) projections. Differential expression analysis was used to annotate clusters into cell types. Fibroblast and immune subclusters were isolated and underwent clustering, differential expression analysis, and UMAP projection to identify different cell state populations present across conditions.

### Statistical analyses

All experiments were powered to identify statistically significant differences and included multiple independent biological replicates. Data were plotted using Prism. Each point represents an individual mouse. For RT-qPCR, three technical replicates per condition were performed. Graphs are mean with s.e.m.

## Acknowledgements

K.J.L. is supported by the Washington University in St. Louis Rheumatic Diseases Research Resource-Based Center (grant no. NIH P30AR073752), the National Institutes of Health (grant nos. R01 HL138466, R01 HL139714, R01 HL151078, R01 HL161185, R35 HL161185), the Leducq Foundation Network (grant no. 20CVD02), the Burroughs Welcome Fund (grant no. 1014782), sponsored research agreement from Amgen, the Children’s Discovery Institute of Washington University and St. Louis Children’s Hospital (grant nos. CH-II-2015-462, CH-II-2017-628, PM-LI-2019-829), the Foundation of Barnes-Jewish Hospital (grant no. 8038-88) and generous gifts from Washington University School of Medicine. S.Y. is supported by the National Institutes of Health (grant no. 5F30HL172583), and the Washington University School of Medicine Medical Scientist Training Program. Schematics used in Figs. 2A, and Extended Data Figs. 1A were created using BioRender (https://BioRender.com). We thank the Genome Technology Access Center at the McDonnell Genome Institute at Washington University School of Medicine for help with genomic analysis. The Center is partially supported by NCI Cancer Center Support Grant no. P30 CA91842 to the Siteman Cancer Center. This publication is solely the responsibility of the authors and does not necessarily represent the official views of the NIH.

## Author contributions

S.Y. and K.J.L. conceived the study and interpreted the data. S.Y. made all the figures. S.Y. and K.J.L. drafted the manuscript. S.Y., A.P., F.K. and A.B. performed all snRNA-seq experiments. S.Y., A.P., An.K. and J.M.A. performed all computational analyses. S.Y. and Y-S.H. performed killing assay in vitro experiments. At.K., C.J.W., and J.M.N. performed in vivo mouse surgeries and echocardiography. S.Y., Y-L.P. and M.O. performed immunohistochemistry and analysed and processed images. S.Y. performed all in vivo experiments. L.Y. performed quantitative PCR experiments. S.D. performed cytokine quantification experiments. M.T., J.W., S.M., K.C. and T.Y. generated and analysed the BiTE reagents. C.B. generated transgenic mice. T.W., B.J. and J.G.R. assisted with experimental design and reagent selection, and provided input for the manuscript. N.H., L.M.W., C-M.L., F.L., N.R., J.A.E., N.S., S.J. and B.A. provided input for the manuscript. All authors contributed to the experimental design, data analysis and interpretation as well as manuscript production. K.J.L. is responsible for all aspects of this manuscript including experimental design, data analysis and manuscript production. All authors approved the final version of the manuscript.

