## Supplemental figures for "Collateral Immune Cell Signaling Compromises the Efficacy of T-cell Engaging Therapies for Cardiac Fibrosis"

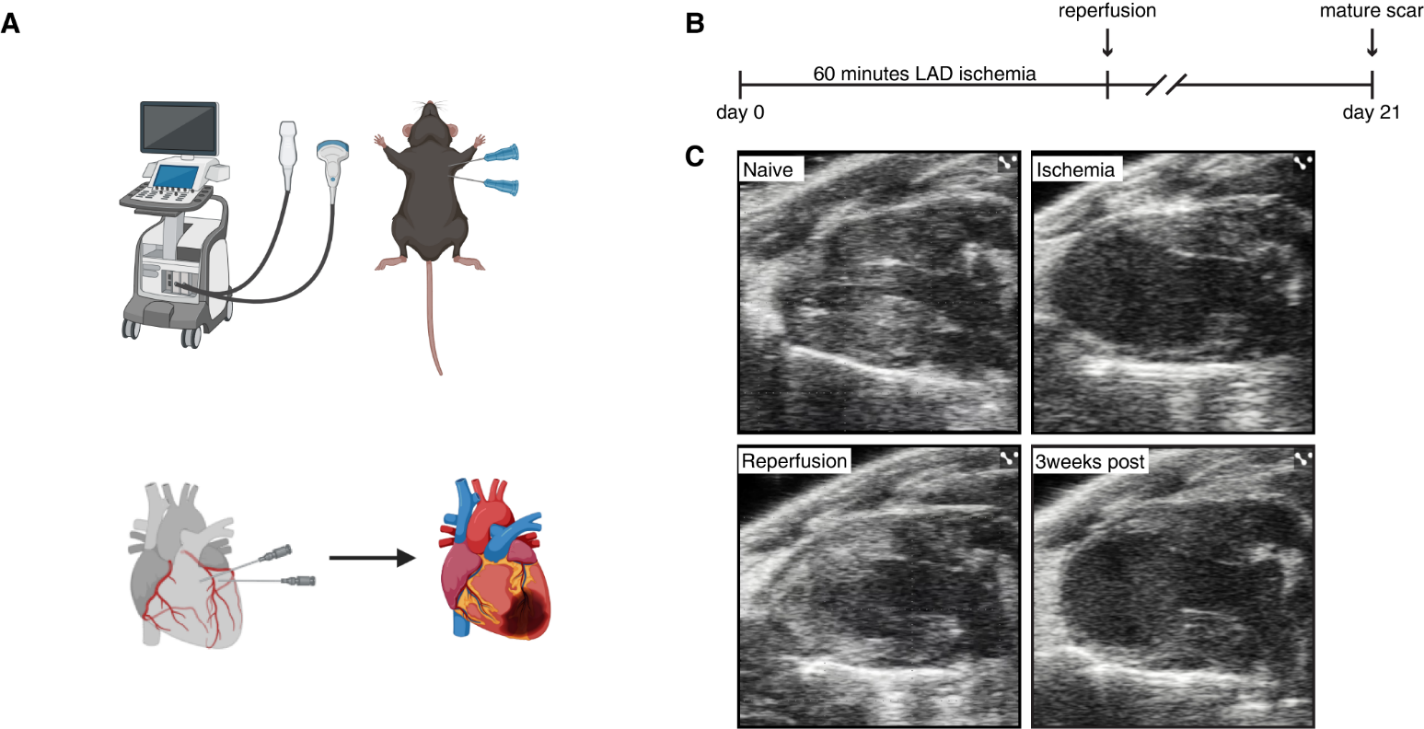

**Supplemental figure 1 – Echo guided IRI.** A) Graphical abstract B) Experimental schematic C) Representative echocardiogram stills before ischemia, during ischemia, during reperfusion, and 3 weeks post IRI

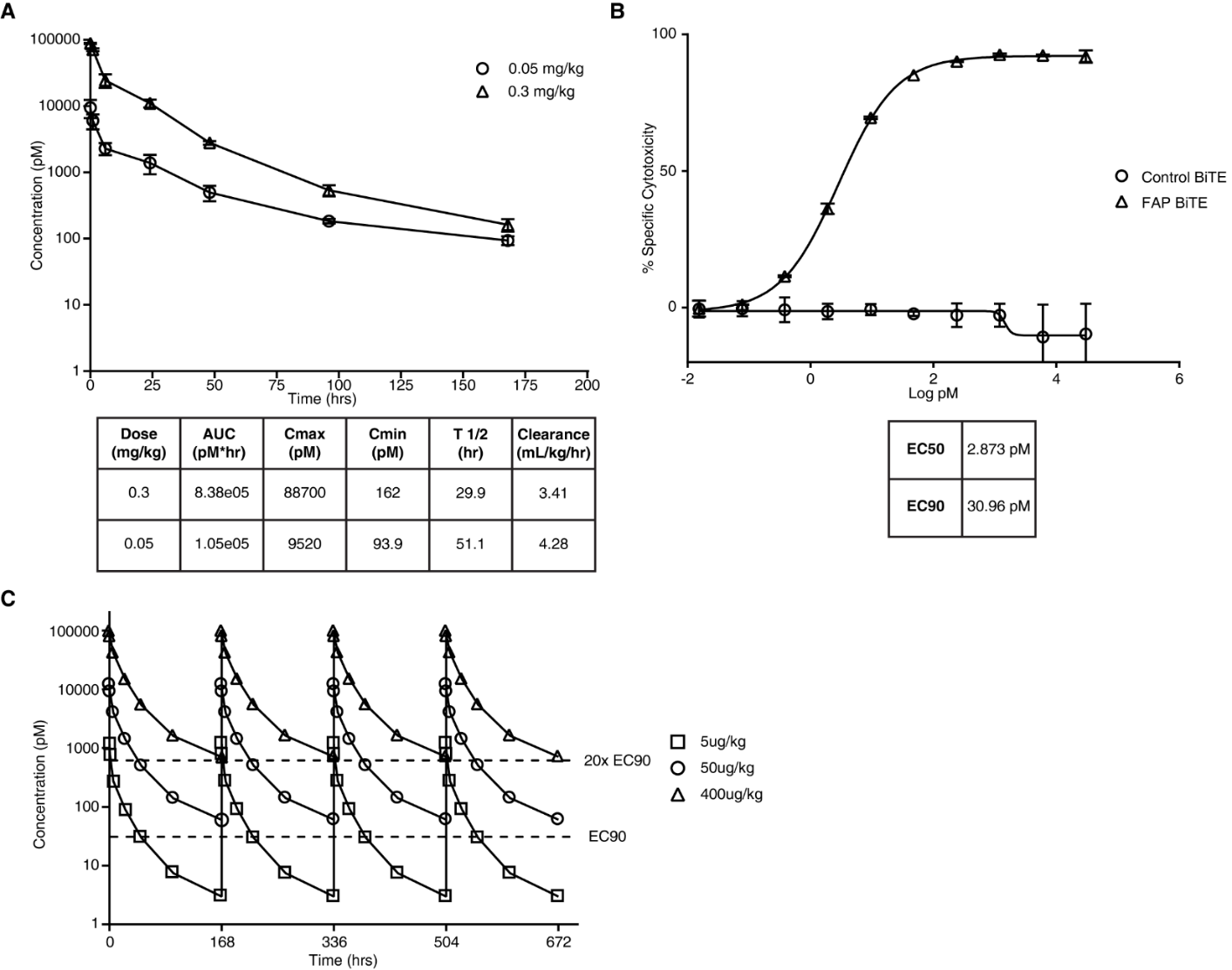

**Supplemental figure 2 – FAP BiTE^®^ molecule pharmacokinetics.** A) FAP BiTE^®^ molecule *in vivo* serum clearance over time with pharmacokinetics B) FAP BiTE^®^ molecule *in vitro* cytotoxicity with EC50 and EC90 C) Simulated FAP BiTE^®^ molecule serum concentration across different weekly doses

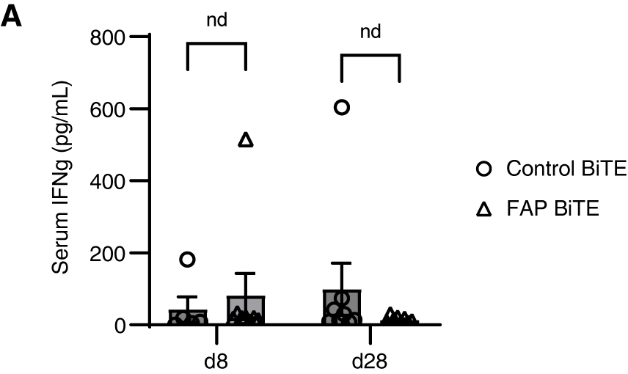

**Supplemental figure 3 – FAP BiTE^®^ molecule does not increase systemic IFNg.** A) Serum IFNg in BiTE^®^ molecule treated hearts day 8 and day 28 after IRI

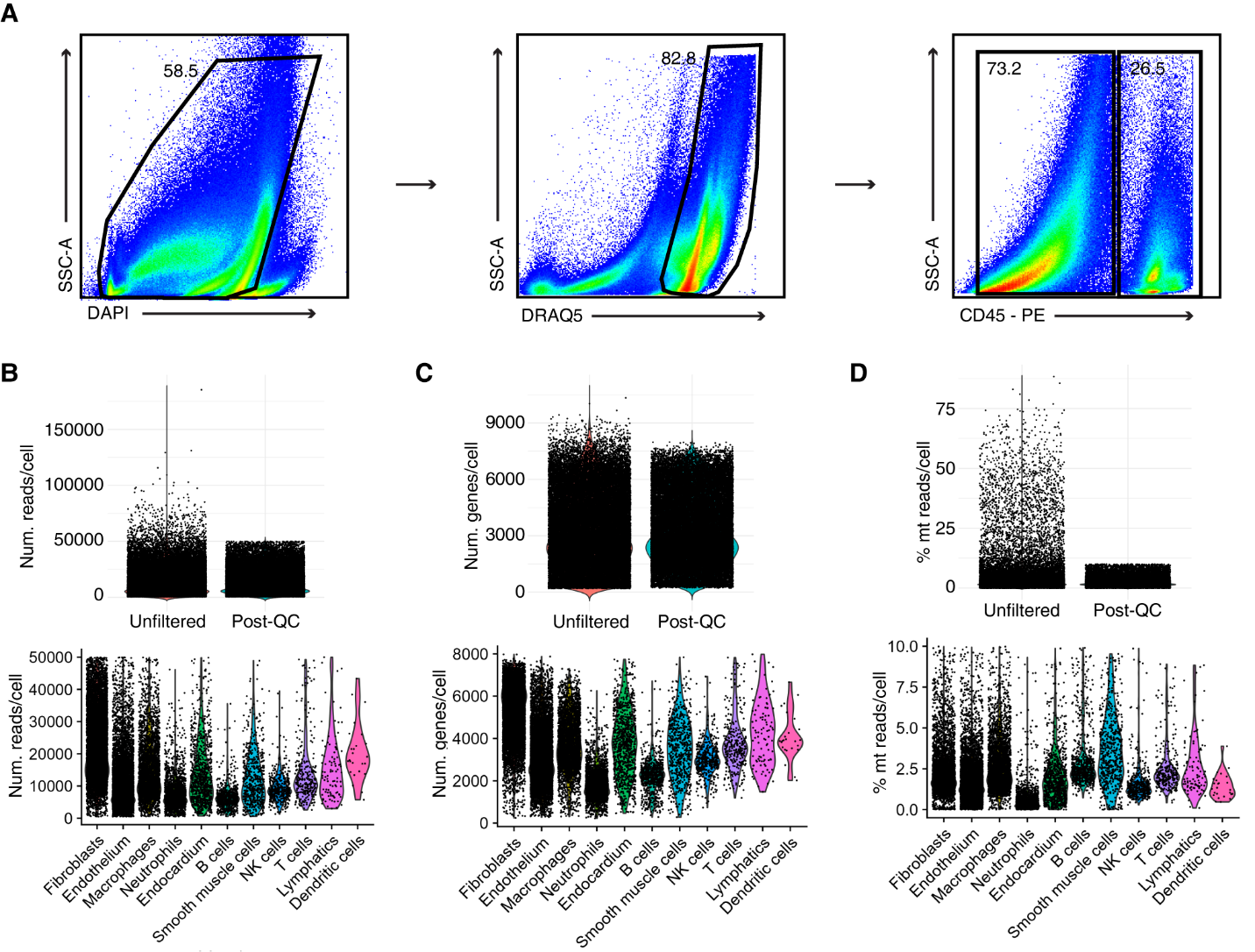

**Supplemental figure 4 – All cell sort and QC metrics.** A) Gating schematic for sort B) Number of reads per cell pre and post QC and across cell types C) Number of genes per cell pre and post QC and across cell types D) Percent mitochondrial reads per cell pre and post QC and across cell types

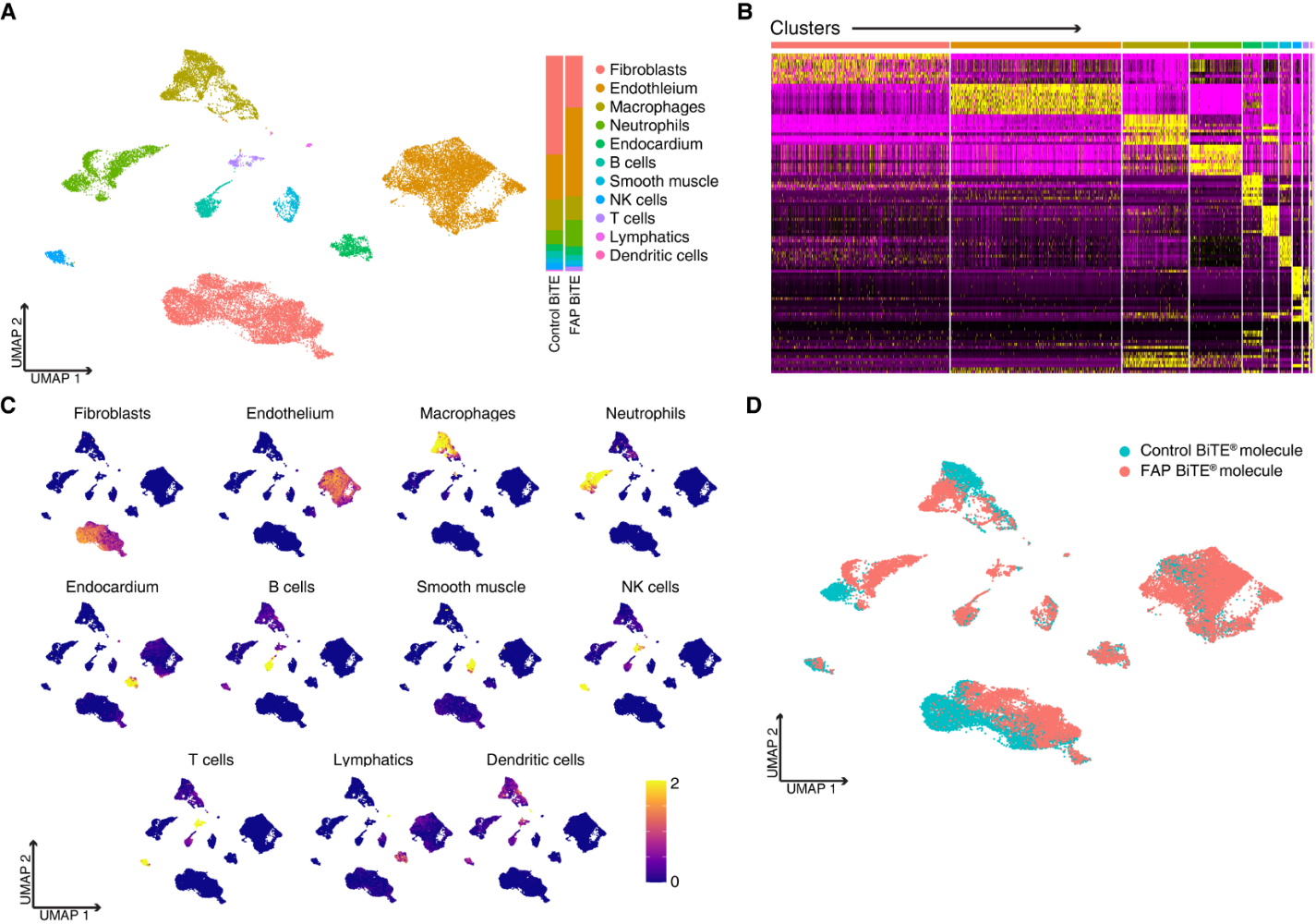

**Supplemental figure 5 – All cell sort UMAP and differentially expressed genes and clustering.** A) UMAP and stack plot split by condition B) Heatmap of differentially expressed genes by cluster C) Plotted z scores of differentially expressed genes by cluster D) Feature plot of cells split by condition

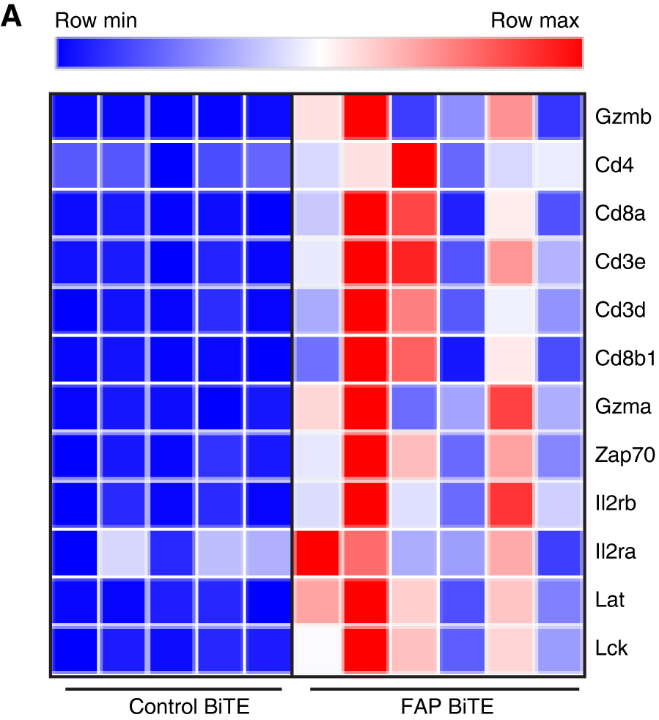

**Supplemental figure 6 – Bulkseq reveals elevated T cell markers in FAP BiTE^®^ molecule treated animals.** A) Heatmap of genes encoding T cell surface markers or downstream signal transducers of T cell receptor signaling

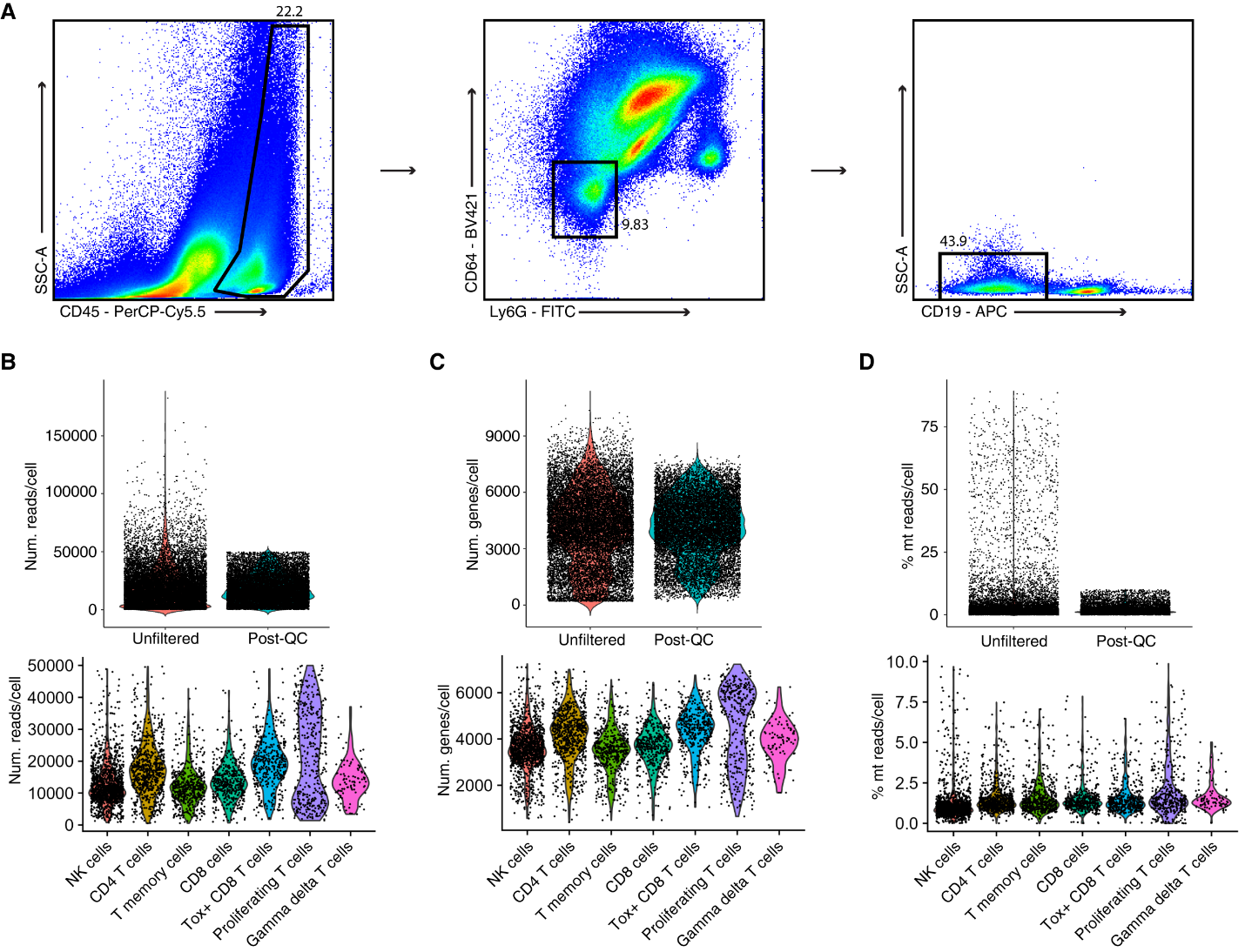

**Supplemental figure 7 – T and NK cell sort and QC metrics.** A) Gating schematic for sort B) Number of reads per cell pre and post QC and across cell types C) Number of genes per cell pre and post QC and across cell types D) Percent mitochondrial reads per cell pre and post QC and across cell types

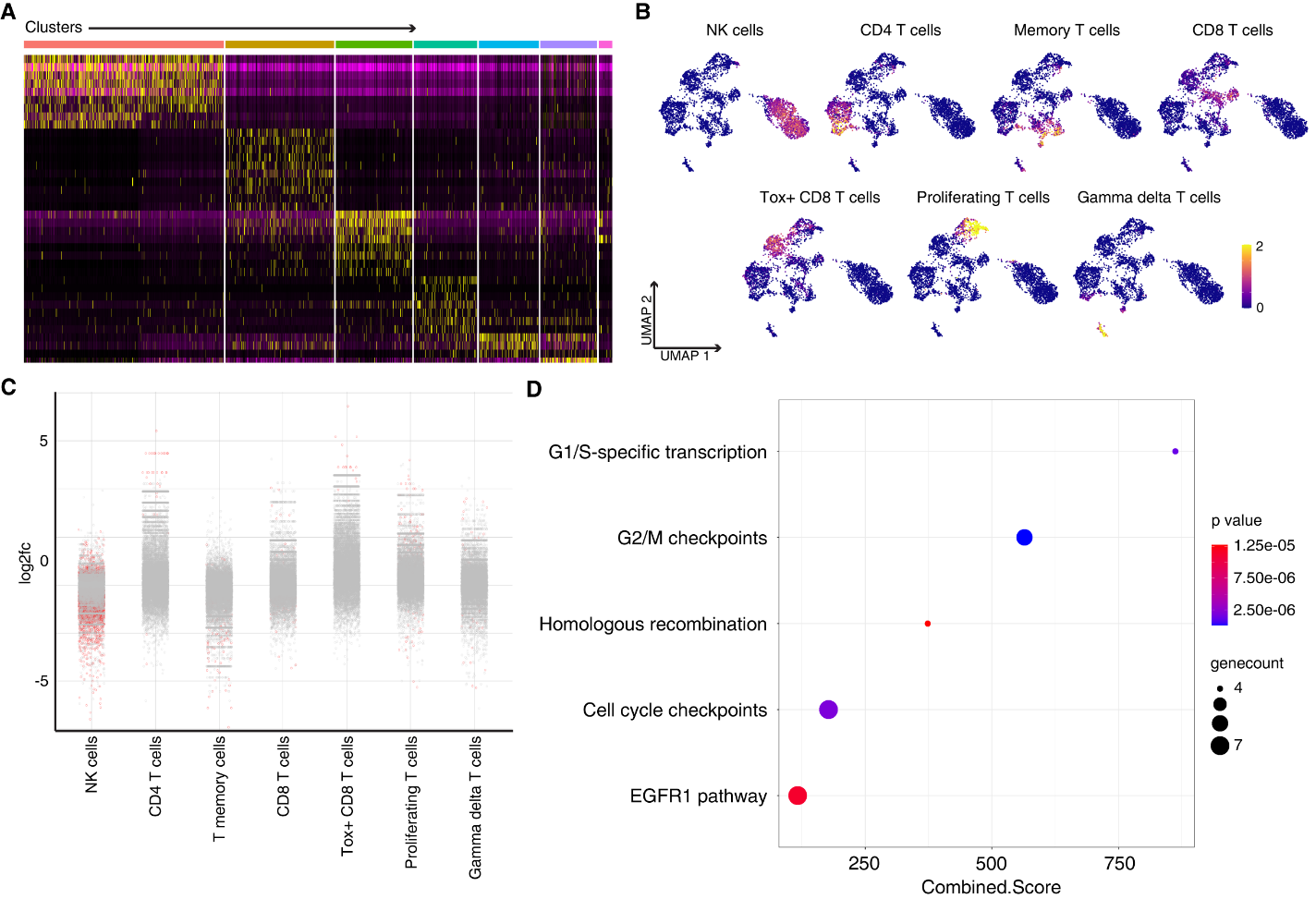

**Supplemental figure 8 – T and NK cell sort differentially expressed genes.** A) Heatmap of differentially expressed genes by cluster B) Plotted z scores of differentially expressed genes by cluster C) Dot plot of differentially expressed genes by cluster D) Pathway analysis of differentially expressed genes by condition

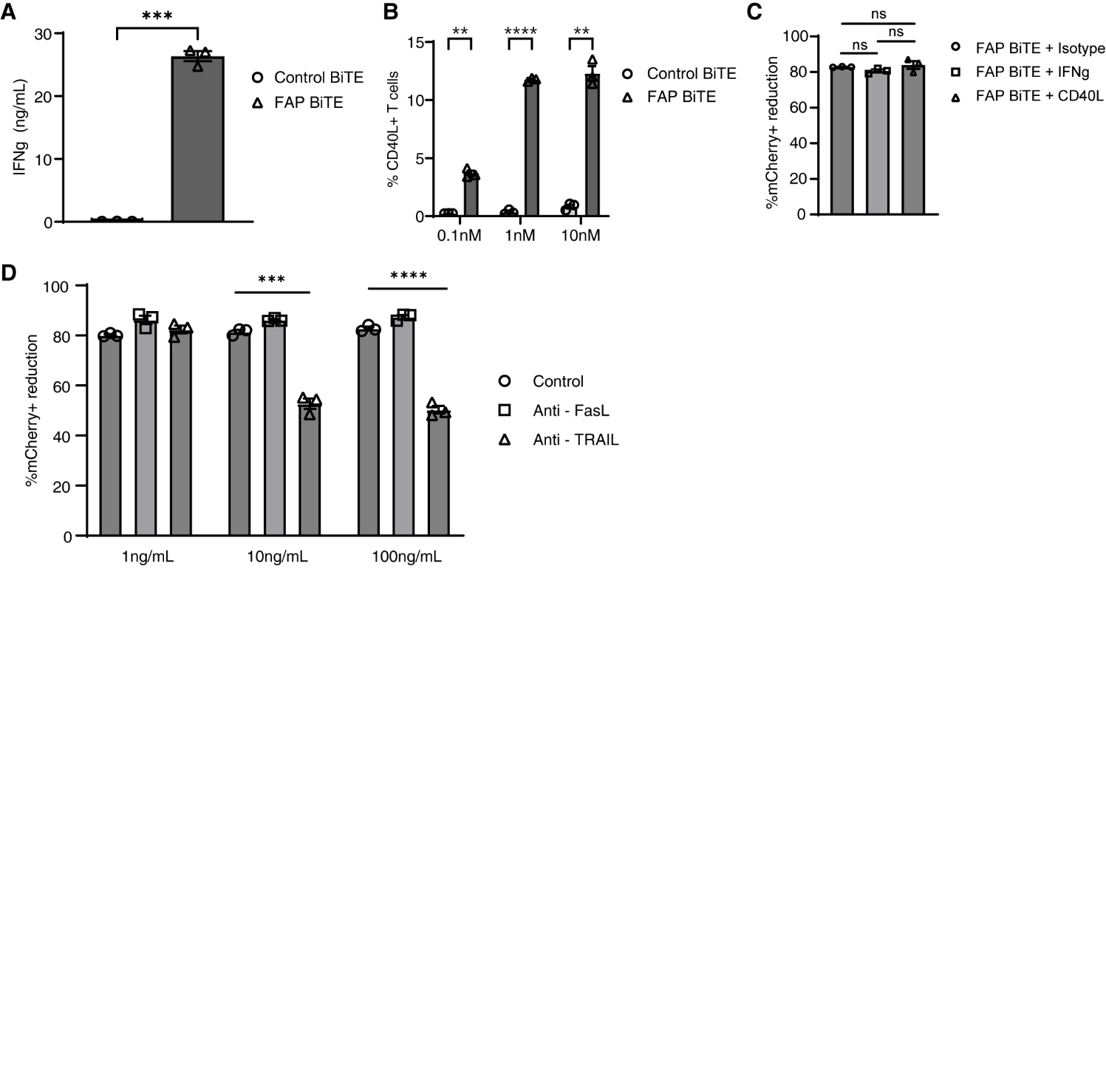

**Supplemental figure 9 – FAP BiTE^®^ molecule mediated killing is independent of IFNg or CD40L signaling.** A) IFNg ELISA of supernatant media collected from 48 hour killing assay coculture B) Percent CD40L positive T cells between conditions across different BiTE^®^ molecule doses C) Killing activity with neutralizing antibodies against IFNg and CD40L D) Killing activity with neutralizing antibodies against FasL and TRAIL

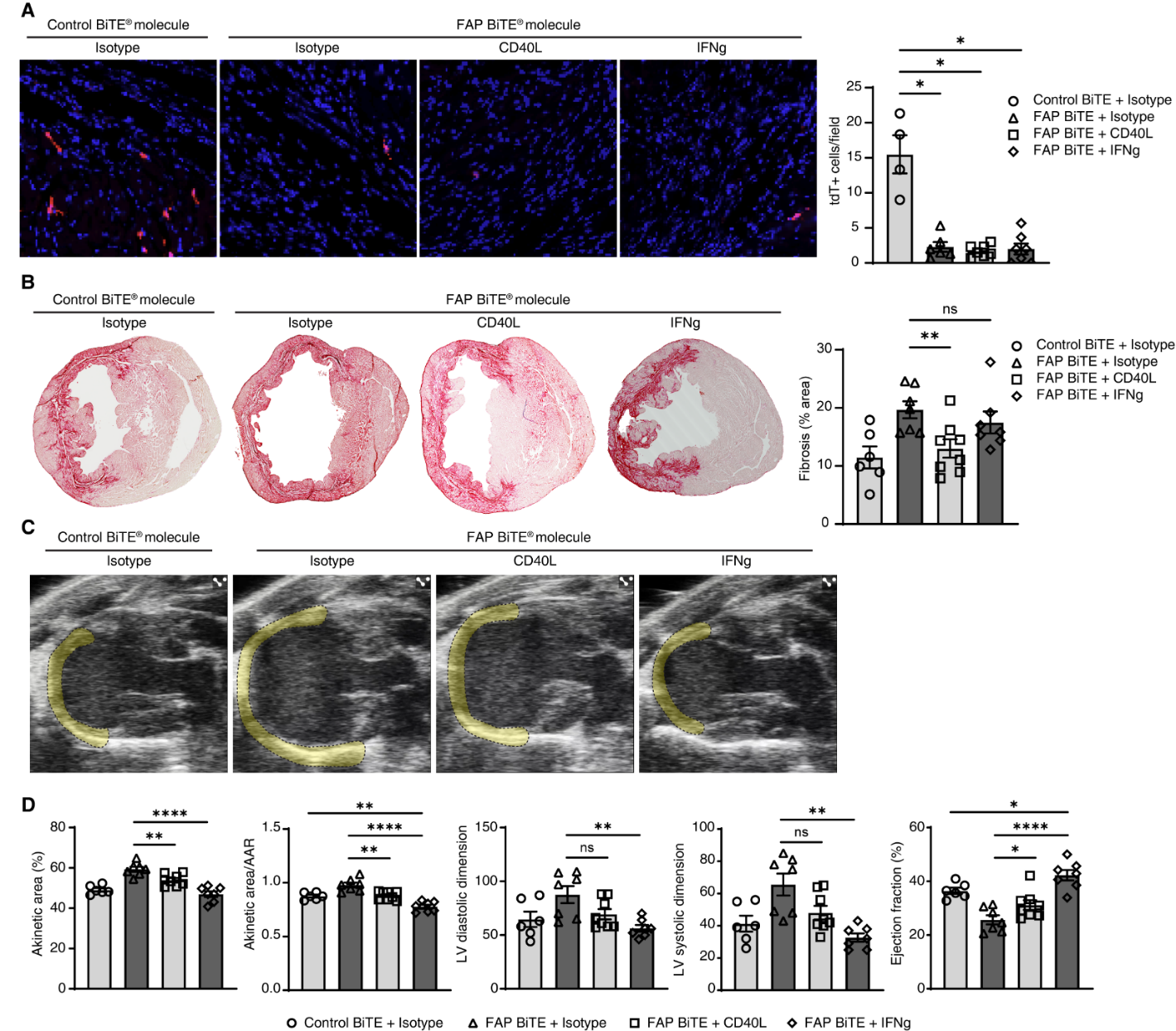

**Supplemental figure 10 – FAP BiTE^®^ molecule rescue with individual neutralization of IFNg or CD40L.** A) tdTomato expression and quantification in FAP BiTE^®^ molecule vs control BiTE^®^ molecule treated mice after IRI with and without IFNg or CD40L neutralization B) Picro Sirius Red staining and quantification C) Representative echocardiogram stills in systole with akinetic area highlighted D) Echocardiogram quantification

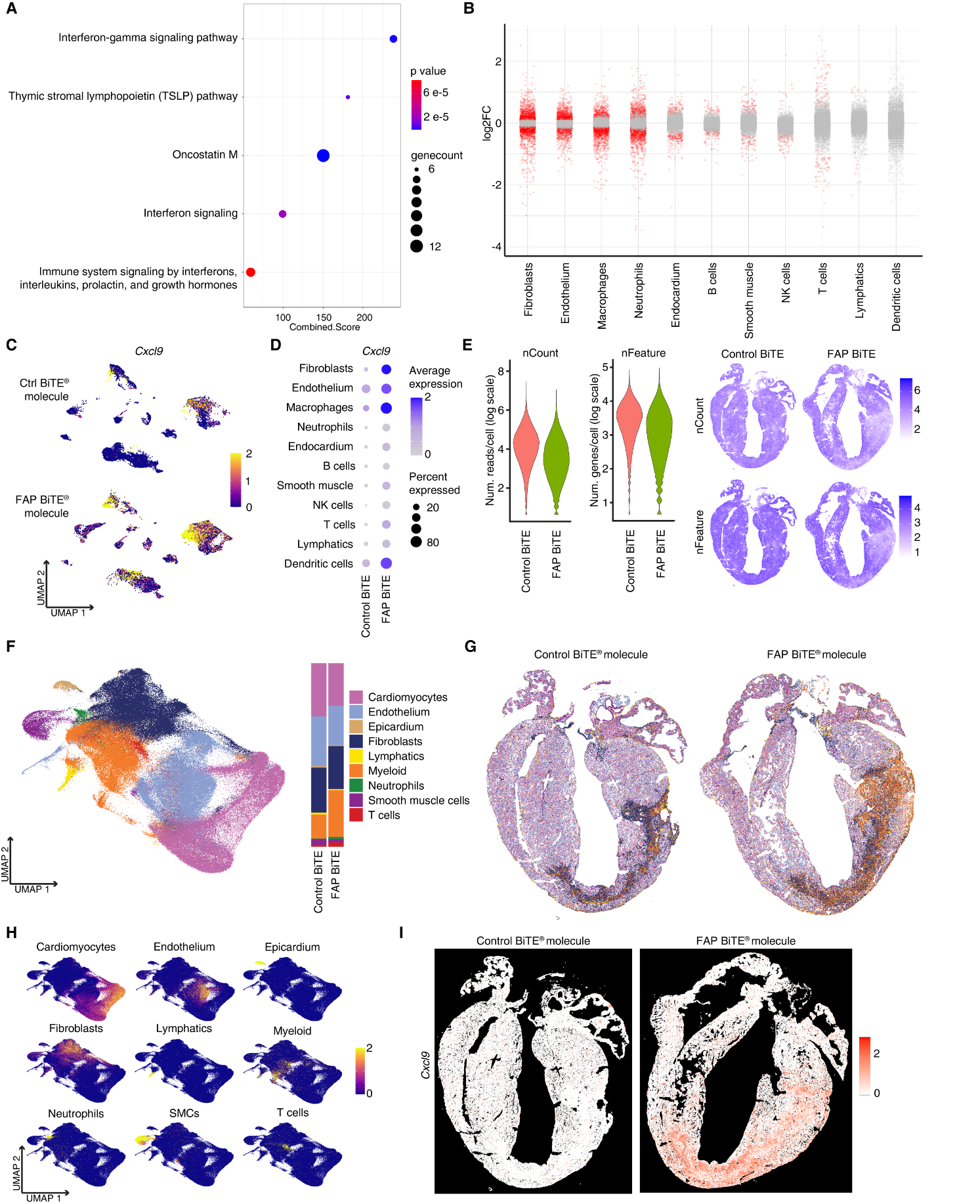

**Supplemental figure 11 – All cell sort scRNAseq and Xenium spatial transcriptomics reveal elevated IFNg signaling in BiTE^®^ molecule treated hearts after IRI.** A) Pathway analysis of differentially expressed genes between control and FAP BiTE^®^ molecule treated hearts after IRI identified with scRNAseq B) Dot plot of differentially expressed genes by cluster C) Feature plot of *Cxcl9* split by condition D) Dot plot of *Cxcl9* split by condition and cluster E) Number of reads and number of genes per cell in Xenium spatial transcriptomics of control vs FAP BiTE^®^ molecule treated hearts day 8 after IRI F) Xenium spatial transcriptomics UMAP and stack plot split by condition G) Spatial UMAP split by condition H) Plotted z scores of differentially expressed genes by cluster I) Spatial feature plot of *Cxcl9* split by condition

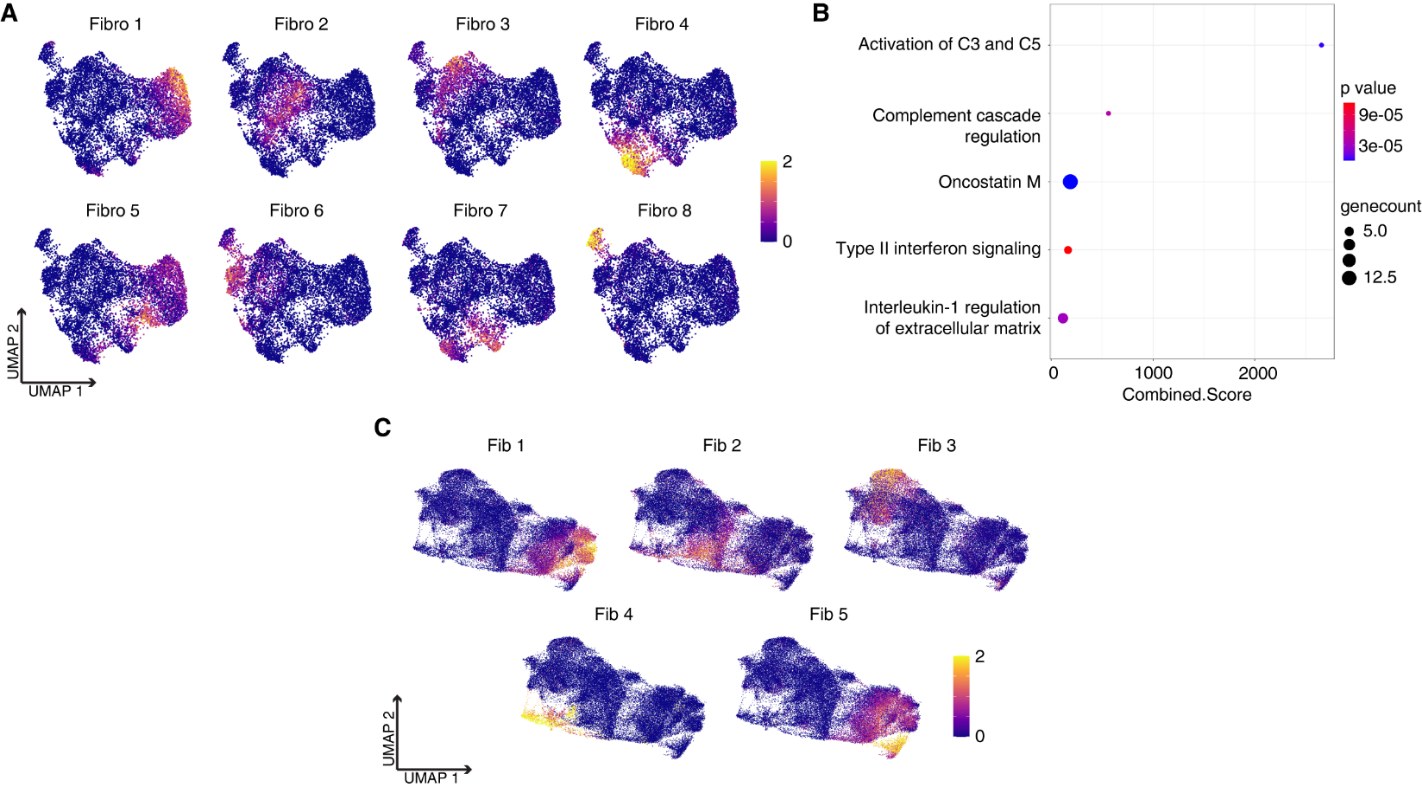

**Supplemental figure 12 – scRNAseq and spatial transcriptomics fibroblast subclustering signature plots and pathway analysis.** A) Plotted z scores of differentially expressed genes by fibroblast subcluster in scRNAseq B) Pathway analysis differentially expressed genes in fibroblasts between conditions in scRNAseq C) Plotted z scores of differentially expressed genes by fibroblast subcluster in spatial transcriptomics

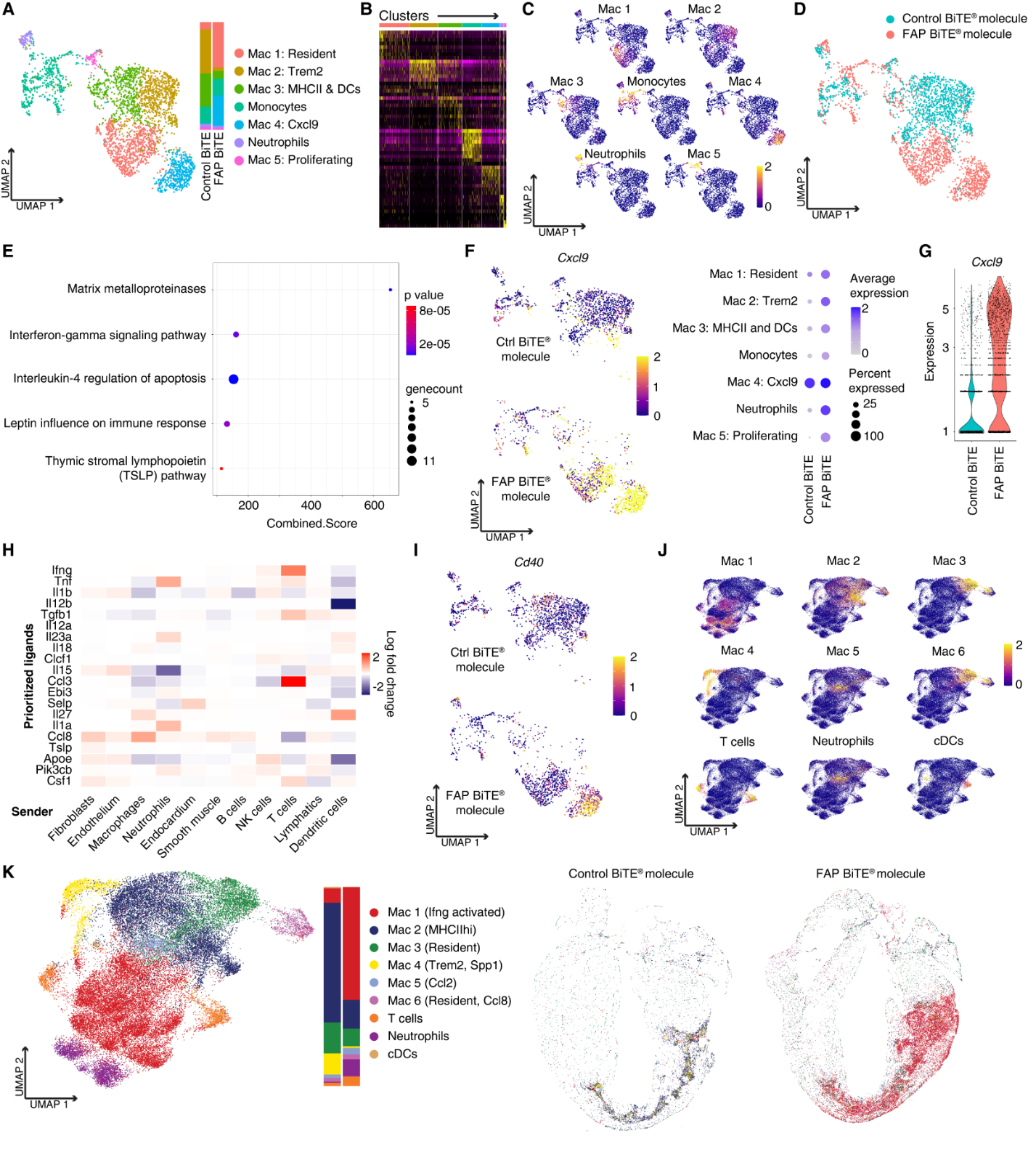

**Supplemental figure 13 – Macrophage subclustering in scRNAseq and spatial transcriptomics**. A) UMAP of subclustered macrophages and stack plot split by condition B) Heatmap of differentially expressed genes by cluster C) Plotted z scores of differentially expressed genes by cluster D) Feature plot of cells split by condition E) Pathway analysis of macrophage subclustering from scRNAseq E) Feature plot and dot plot of *Cxcl9* split by condition and subcluster G) Violin plot of *Cxcl9* split by condition H) NicheNet receptor ligand analysis heatmap with macrophage as receiver I) Feature plot of *Cd40* split by condition J) Plotted z scores of differentially expressed genes by cluster in macrophages with spatial transcriptomics K) Spatial transcriptomic UMAP of macrophage subclustering with stack plot split by condition
